# Mesenchymal Stromal Cell Senescence Induced by *Dnmt3a*-Mutant Hematopoietic Cells is a Targetable Mechanism Driving Clonal Hematopoiesis and Initiation of Hematologic Malignancy

**DOI:** 10.1101/2024.03.28.587254

**Authors:** Jayna J. Mistry, Kira A. Young, Patricia A. Colom Díaz, Inés Fernández Maestre, Ross L. Levine, Jennifer J. Trowbridge

## Abstract

Clonal hematopoiesis (CH) can predispose to blood cancers due to enhanced fitness of mutant hematopoietic stem and progenitor cells (HSPCs), but the mechanisms driving this progression are not understood. We hypothesized that malignant progression is related to microenvironment-remodelling properties of CH-mutant HSPCs. Single-cell transcriptomic profiling of the bone marrow microenvironment in *Dnmt3a*^R878H/+^ mice revealed signatures of cellular senescence in mesenchymal stromal cells (MSCs). *Dnmt3a*^R878H/+^ HSPCs caused MSCs to upregulate the senescence markers SA-β-gal, BCL-2, BCL-xL, *Cdkn1a* (p21) and *Cdkn2a* (p16), *ex vivo* and *in vivo*. This effect was cell contact-independent and can be replicated by IL-6 or TNFα, which are produced by *Dnmt3a*^R878H/+^ HSPCs. Depletion of senescent MSCs *in vivo* reduced the fitness of *Dnmt3a*^R878H/+^ hematopoietic cells and the progression of CH to myeloid neoplasms using a sequentially inducible *Dnmt3a*;*Npm1*-mutant model. Thus, *Dnmt3a*-mutant HSPCs reprogram their microenvironment via senescence induction, creating a self-reinforcing niche favoring fitness and malignant progression.

**Statement of Significance:** Mesenchymal stromal cell senescence induced by *Dnmt3a*-mutant hematopoietic stem and progenitor cells drives clonal hematopoiesis and initiation of hematologic malignancy.

## Introduction

Aging is associated with functional decline of the hematopoietic system(*1*) and development of clonal hematopoiesis (CH)(*2*). CH is defined by over-representation of hematopoietic stem cells (HSCs) and their progeny carrying somatic mutations or variants that confer a selective advantage. While CH is a benign condition, several of the somatic mutations found in CH are associated with hematologic malignancies including myeloid neoplasms, such as mutations in the DNA methyltransferase *DNMT3A*(*3*). The mechanisms by which CH progresses into myeloid neoplasms and other hematologic malignancies remain incompletely understood. A recent clonal hematopoiesis risk score paradigm defined risk of myeloid neoplasms (myelodysplastic syndrome, myeloproliferative neoplasms, acute myeloid leukemia (AML), and overlap conditions) in individuals with CH based on their age, clone size, genotype, and complete blood count parameters(*4*). However, to understand the functional mechanisms underlying risk of hematologic malignancies, and to develop intervention strategies, prospective experimental data are needed.

In the context of hematologic malignancies, non-hematopoietic cell types in the bone marrow microenvironment have emerged as a key partner in creating and maintaining a malignant niche. The bone marrow microenvironment undergoes extensive remodeling in malignancy which contributes to disease development(*5-8*). For example, secretion of vascular endothelial growth factor (VEGF) by malignant hematopoietic cells stimulates angiogenesis from endothelial cells, which promotes malignant cell survival and proliferation(*5-8*). Bone marrow mesenchymal stromal cells (MSCs) isolated from patients with acute myeloid leukemia (AML) have increased adipogenic potential(*9*).

Lipolysis of BM adipocytes by leukemic stem cells(*10*) and leukemic blasts(*11*) fuel fatty acid oxidation and oxidative phosphorylation, which the leukemic stem cells(*12*) and leukemic blasts(*11*) depend upon for survival. AML blasts have also been shown to increase oxidative stress, which increases mitochondrial biogenesis in MSCs(*13, 14*).

This is advantageous to AML growth as the mitochondria in MSCs can be acquired by blast cells via production of tunneling nanotubes(*13, 14*). In primary patient samples from individuals with myelodysplastic syndrome, multiple myeloma, myeloproliferative neoplasms and AML, malignant hematopoietic cells have been demonstrated to induce cellular senescence of MSCs(*15-18*). Senescence is a physiological program characterized by irreversible cell cycle arrest, altered gene expression, and secretion of numerous pro-inflammatory cytokines, chemokines, growth factors, and proteases known as the senescence-associated secretory phenotype (SASP). Given that selective elimination of senescent MSCs *in vivo* extends survival in a murine AML model(*18*), targeting senescent cells in the BM microenvironment has emerged as an attractive target for therapeutic intervention in myeloid neoplasms.

In comparison to the body of knowledge regarding the role of non-hematopoietic cell types in maintaining a niche for malignant cells, little is known regarding the extent to which mutant clones in CH shape the BM microenvironment. Furthermore, it remains unknown whether therapeutic strategies to intercept clonal selection and expansion in the context of CH and pre-malignancy are effective in prevention or delay of myeloid neoplasms. Here, we report altered characteristics of the BM microenvironment in a model of *DNMT3A*-mutant CH, including induction of MSC senescence. We show that induction of MSC senescence by *Dnmt3a*-mutant CH is therapeutically targetable. Depletion of senescent cells not only reduces the competitive advantage of CH-mutant cells but also reduces the progression from CH to myeloid neoplasms.

## Results

### *Dnmt3a*-Mutant Hematopoiesis Induces Transcriptional Signatures of Senescence in Bone Marrow Mesenchymal Stromal Cells

Our group recently reported that a middle-aged BM microenvironment contributes to HSC aging phenotypes(*19*), and *Dnmt3a*^R878H/+^ HSCs exhibit increased selective advantage over wild-type HSCs as well as altered lineage output(*20*). To determine the cellular and molecular effects of *Dnmt3a*^R878H/+^ hematopoietic cells on non-hematopoietic cells of the aged BM microenvironment, we transplanted CD45.2^+^ BM cells from *Dnmt3a*^R878H/+^ Fgd5-Cre LSL-tdTomato or control Fgd5-Cre LSL-tdTomato mice into wild-type middle-aged CD45.1^+^ recipient mice following busulfan conditioning. At 12 weeks post-transplant, we harvested BM and performed single cell RNA-sequencing (scRNA-seq) on enriched hematopoietic and non-hematopoietic cell fractions using established protocols(*21*) (*n* = 4 biological replicates per condition) (Figure 1A). Cells were sorted based on the following cell surface marker combinations: mature hematopoietic cells (CD45.2^+^ tdTomato^+^ Lineage^+^), committed hematopoietic progenitor cells (CD45.2^+^ tdTomato^+^ Lineage^-^ c-Kit^+^ Sca-1^-^), hematopoietic stem and multipotent progenitor cells (CD45^+^ tdTomato^+^ Lineage^-^ c-Kit^+^ Sca-1^+^), endothelial cells (CD45^-^ Ter119^-^ CD41^-^ CD31^+^), and mesenchymal stromal cells (CD45^-^ Ter119^-^ CD41^-^ CD31^-^ CD71^-^ CD51^+^) (Figure 1A). A total of 61,105 cells were sequenced and we identified a total of 26 clusters including the hematopoietic cell types hematopoietic stem cell/multipotent progenitor (HSC/MPP), granulocyte-macrophage-primed multipotent progenitor (MPP^G/M^), lymphoid-primed multipotent progenitor (MPP^Ly^), megakaryocyte-erythroid progenitor (MEP), erythroid progenitor (EryPro), erythroblast (Ery), mast-eosinophil-basophil progenitor (MEoBasoPro), cycling progenitor (Cycling), granulocyte-macrophage progenitor (GMP), neutrophil progenitor (NeutPro), immature neutrophil (ImmNeut), neutrophil (Neut), monocyte progenitor (MonoPro), monocyte (Mono), macrophage (Mac), monocyte/dendritic cell (Mono/DC), dendritic cell (DC), pre-B cell (PreB), pro-B cell (ProB), B cell (B) and natural killer/T cells (NK/T), and the non-hematopoietic cell types adipocyte-lineage-primed mesenchymal stromal cells (MSC-Adipo), osteo-lineage-primed mesenchymal stromal cells (MSC-Osteo), osteoblast (Osteo), arteriolar endothelial cell (EC-Arteriolar), and sinusoidal endothelial cell (EC-Sinusoidal) (Figure 1B). We observed an increased frequency of *Dnmt3a*^R878H/+^ HSC/MPP and NeutPro compared to control, and a decreased frequency of *Dnmt3a*^R878H/+^ MPP^Ly^ compared to control (Supplemental Figure 1A). Of the genes upregulated in *Dnmt3a*^R878H/+^ vs. control HSC/MPPs, enrichment of transcriptional signatures of leukocyte migration, neutrophil activation, immune system processes, and myeloid differentiation were observed (Supplemental Figure 1B). Of the genes downregulated in *Dnmt3a*^R878H/+^ vs. control HSC/MPPs, enrichment of transcriptional signatures of type II interferon response and cytokine response were observed (Supplemental Figure 1B), consistent with previous literature demonstrating that loss of *Dnmt3a* results in reduced interferon responsiveness(*22, 23*).

**Figure 1.**
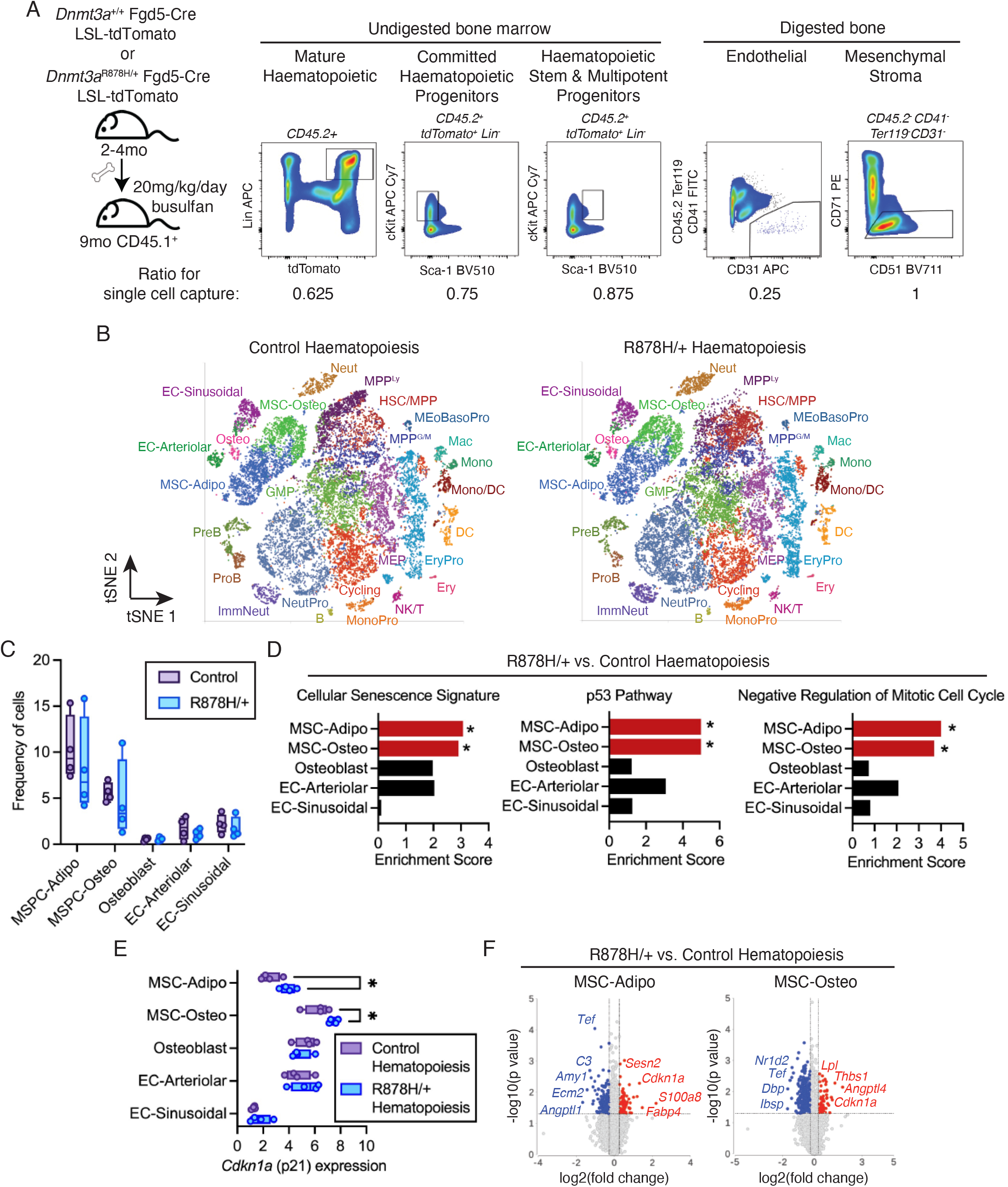
*Dnmt3a*-mutant hematopoiesis is associated with transcriptional signatures of senescence in MSCs. (**A**) Schematic of experimental design for single-cell RNA-seq (scRNA-seq) (*n* = 4 replicates per condition). Representative flow cytometry plots showing cell populations and cell ratios sorted for scRNA-seq. (**B**) t-SNE plots showing 22 annotated cell clusters. B: B cells, DC: dendritic cells, EC-arteriolar: arteriolar endothelial cells, EC-sinusoidal: sinusoidal endothelial cells, Ery: erythrocytes, EryPro: erythroid progenitor cells, GMP: granulocyte-macrophage progenitor cells, HSC/MPP: hematopoietic stem and multipotent progenitor cells, ImmNeut: immature neutrophils, Mac: macrophages, MEoBasoPro: mast-eosinophil-basophil progenitor cells, MEP: megakaryocyte-erythroid progenitor cells, Mono: monocytes, MonoPro: monocyte progenitor cells, MPP^G/M^: granulocyte-macrophage-primed multipotent progenitor cells, MPP^Ly^: lymphoid-primed multipotent progenitor cells, MSC-Adipo: adipocyte-primed mesenchymal stromal cells, MSC-Osteo: osteo-primed mesenchymal stromal cells, Neut: neutrophils, NeutPro: neutrophil progenitor cells, NK/T: natural killer and T cells, Osteo: osteoblasts, PreB: pre-B cells, ProB: pro-B cells. (**C**) Frequency of non-hematopoietic cell types in mice with control or *Dnmt3a*^R878H/+^ hematopoiesis. *n* = 4. (**D**) Enrichment of cellular senescence, p53 pathway and negative regulation of mitotic cell cycle gene signatures in non-hematopoietic cell types in mice with *Dnmt3a*^R878H/+^ vs. control hematopoiesis. \**P* < 0.05 by gene set enrichment analysis. (**E**) *Cdkn1a* expression in non-hematopoietic cell types in mice with control or *Dnmt3a*^R878H/+^ hematopoiesis. \**P* < 0.05 by two-way ANOVA with uncorrected Fisher’s LSD. (**F**) Volcano plots showing significantly differentially expressed genes (increased: red, decreased: blue) in MSC-Adipo and MSC-Osteo clusters in mice with *Dnmt3a*^R878H/+^ vs. control hematopoiesis.

We then evaluated non-hematopoietic cells in the BM microenvironment. We did not observe differences in the proportions of non-hematopoietic cell populations in *Dnmt3a*^R878H/+^ compared to control bone marrow (Figure 1C). However, there were significantly differentially expressed genes when comparing non-hematopoietic cell populations isolated from mice with *Dnmt3a*^R878H/+^ vs. control hematopoietic cells.

Among the most enriched signatures in MSC-Osteo and MSC-Adipo populations exposed to *Dnmt3a*^R878H/+^ vs. control hematopoiesis was a cellular senescence gene signature (Figure 1D). This gene signature was uniquely enriched in MSCs and not observed in osteoblasts or ECs exposed to *Dnmt3a*^R878H/+^ vs. control hematopoiesis (Figure 1D). MSC-Osteo and MSC-Adipo populations exposed to *Dnmt3a*^R878H/+^ hematopoiesis also exhibited increased expression of genes enriched for the p53 pathway and negative regulation of mitotic cell cycle (Figure 1D), both of which are associated with replicative senescence. This program included increased expression of the canonical senescence-associated gene *Cdkn1a* (p21) (Figure 1E-F). Together, these data show that *Dnmt3a*^R878H/+^ hematopoiesis is associated with molecular signatures of senescence occurring selectively within MSCs in the BM microenvironment.

### *Dnmt3a*-Mutant Hematopoiesis Induces Molecular Markers of MSC Senescence

We next assessed the cellular phenotypes associated with molecular signatures of senescence. To determine the extent to which Dnmt3a^R878H/+^ hematopoietic cells initiate MSC senescence, we transplanted control or *Dnmt3a*^R878H/+^ BM into wild-type young CD45.1^+^ recipient mice following busulfan conditioning (Figure 2A). At 12 weeks, we harvested peripheral blood, BM hematopoietic cells and BM non-hematopoietic cells.

**Figure 2.**
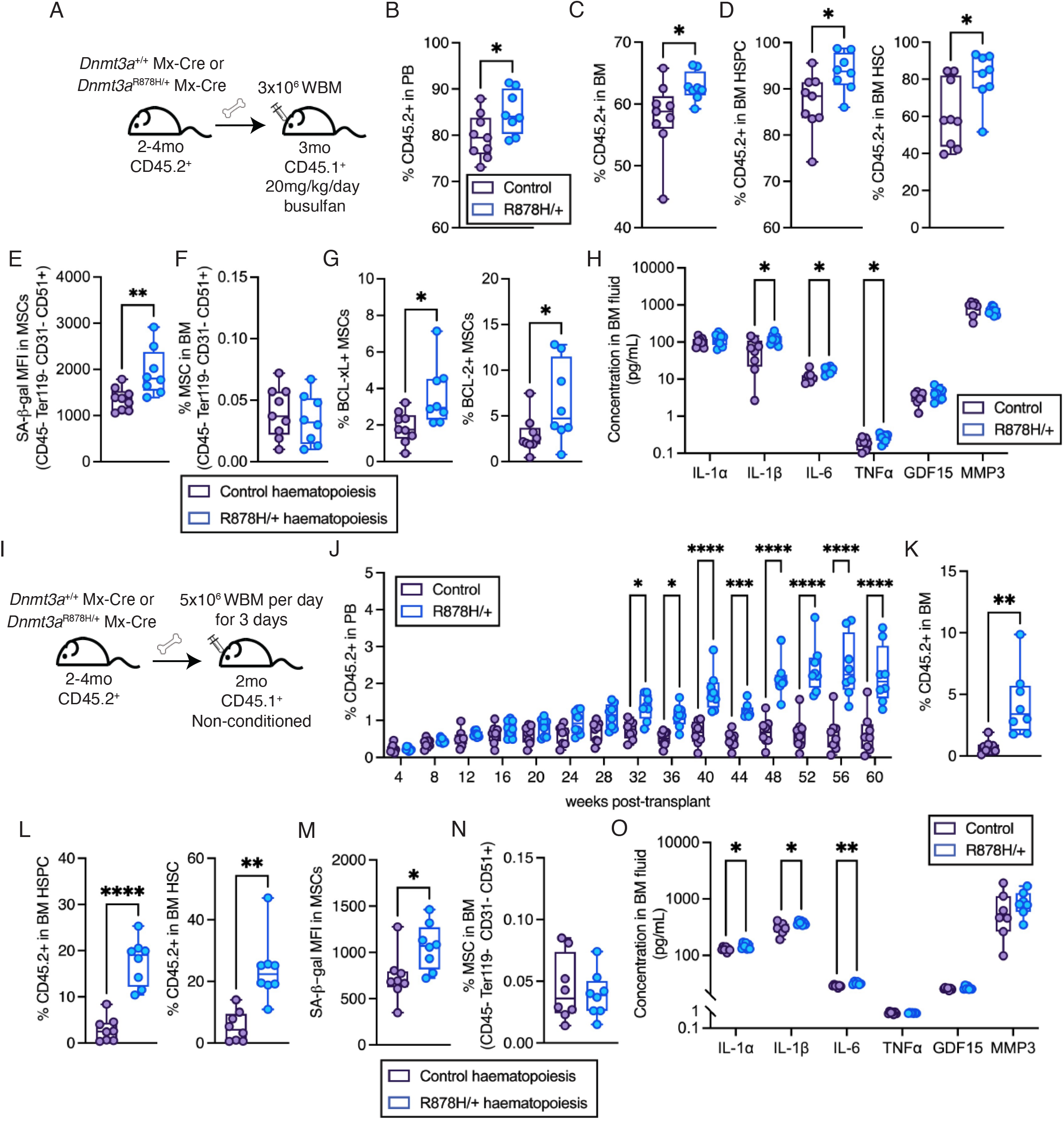
*Dnmt3a*-mutant hematopoiesis induces MSC senescence *in vivo*. (**A**) Schematic of busulfan-conditioned transplant design. (**B-D**) Frequency of donor (CD45.2^+^) cells in (**B**) peripheral blood (PB), (**C**) bone marrow (BM), (**D**) hematopoietic stem and progenitor cells (HSPC) and hematopoietic stem cells (HSC) of mice at 12 weeks post-transplant. \**P* < 0.05 by Mann-Whitney test (*n* = 8-9). (**E**) Mean fluorescence intensity (MFI) of SA-β-gal in MSCs isolated from mice with control or *Dnmt3a*^R878H/+^ hematopoiesis. \*\**P* < 0.01 by Mann-Whitney test (*n* = 8-9). (**F**) Frequency of MSCs in the bone marrow of mice with control or *Dnmt3a*^R878H/+^ hematopoiesis (*n* = 8-9). (**G**) Frequency of BCL-xL^+^ and BCL-2^+^ MSCs in the bone marrow of mice with control or *Dnmt3a*^R878H/+^ hematopoiesis. \**P* < 0.05 by Mann-Whitney test (*n* = 8-9). (**H**) Concentration of cytokines and growth factors in bone marrow fluid of mice with control or *Dnmt3a*^R878H/+^ hematopoiesis assessed by multiplexed ELISA. \**P* < 0.05 by Mann-Whitney test (*n* = 6-9). (**I**) Schematic of non-conditioned transplant design. (**J**) Frequency of donor cells in PB of recipient mice. \**P* < 0.05, \*\*\**P* < 0.001, \*\*\*\**P* < 0.0001 by two-way ANOVA with Sidak’s multiple comparisons test (*n* = 8). (**K-L**) Frequency of donor cells in (**K**) BM and (**L**) HSPC and HSC of recipient mice at 60 weeks post-transplant. \*\**P* < 0.01, \*\*\*\**P* < 0.0001 by Mann-Whitney test (*n* = 8). (**M**) MFI of SA-β-gal in MSCs isolated from mice with control or *Dnmt3a*^R878H/+^ hematopoiesis. \**P* < 0.05 by Mann-Whitney test (*n* = 8). (**N**) Frequency of MSCs in the bone marrow of mice with control or *Dnmt3a*^R878H/+^ hematopoiesis (*n* = 8). (**O**) Concentration of cytokines and growth factors in bone marrow fluid of mice with control or *Dnmt3a*^R878H/+^ hematopoiesis assessed by multiplexed ELISA. \**P* < 0.05 by Mann-Whitney test (*n* = 7).

The gating strategy for hematopoietic stem and progenitor cells (HSPCs) is shown in Supplemental Figure 2A. As expected, *Dnmt3a*^R878H/+^ hematopoietic cells had increased engraftment in the peripheral blood (Figure 2B, Supplemental Figure 2BC) and bone marrow (Figure 2C) and had higher proportions of hematopoietic stem and progenitor cells (HSPCs) and HSCs compared to control transplant mice (Figure 2D). Using flow cytometry, increased intracellular senescence-associated beta-galactosidase (SA-β-gal) was observed in MSCs (CD45^-^ Ter119^-^ CD31^-^ PDGFRα^-^ CD51^+^) in mice with *Dnmt3a*^R878H/+^ vs. control hematopoiesis (Figure 2E), without a change in the frequency of MSCs (Figure 2F). Intracellular protein expression of the senescence-associated anti-apoptotic proteins BCL-xL and BCL-2 were also increased in MSCs in mice with *Dnmt3a*^R878H/+^ vs. control hematopoiesis (Figure 2G). To examine the physiological consequences of MSC senescence, we assessed SASP by quantitation of IL-1α, IL-1β, IL-6, TNFα, GDF15, and MMP3 in the BM fluid. Mice transplanted with *Dnmt3a*^R878H/+^ hematopoiesis exhibited elevated levels of IL-1β, IL-6 and TNFα in the BM fluid compared to control (Figure 2H). Consistent with our transcriptional data, there was no change in SA-β-gal within or frequency of endothelial cells (ECs) (CD45^-^ Ter119^-^ CD71^-^ CD31^+^) in mice with *Dnmt3a*^R878H/+^ vs. control hematopoiesis (Supplemental Figure 2D), further demonstrating that senescence occurs selectively within MSCs in the BM microenvironment.

Our experimental transplantation designs employ conditioning methods to enable reconstitution of *Dnmt3a*^R878H/+^ hematopoiesis in the context of a wild-type BM microenvironment. However, conditioning methods including ionizing radiation and busulfan have been shown to induce some degree of senescence in non-hematopoietic and hematopoietic BM cells(*24, 25*). To avoid this confounding factor and more closely model human clonal hematopoiesis, we transplanted *Dnmt3a*^R878H/+^ or control BM hematopoietic cells into wild-type young CD45.1^+^ recipient mice in the absence of conditioning (Figure 2I). This resulted in a small amount of detectable engraftment (0.2-0.5%) within the peripheral blood at 4 weeks post-transplant (Figure 2J). Over 60 weeks, *Dnmt3a*^R878H/+^ hematopoietic cells increased their long-term multilineage reconstitution up to 4%, significantly higher than control hematopoietic cells which showed stable low-level engraftment (Figure 2J). At 60 weeks post-transplant, *Dnmt3a*^R878H/+^ hematopoietic cells had increased engraftment in the bone marrow (Figure 2K) and had higher proportions of HSPCs and HSCs compared to control transplant mice (Figure 2L). In addition, *Dnmt3a*^R878H/+^ peripheral blood had increased proportions of myeloid cells and granulocytes compared to control (Supplemental Figure 2E-F). Increased intracellular SA-β-gal was observed in MSCs in mice with *Dnmt3a*^R878H/+^ vs. control hematopoiesis (Figure 2M), without a change in the frequency of MSCs (Figure 2N). To examine the physiological consequences of MSC senescence, we assessed SASP. Mice transplanted with *Dnmt3a*^R878H/+^ hematopoiesis exhibited elevated levels of IL-1α, IL-1β and IL-6 in the BM fluid compared to control (Figure 2O). There was no change in SA-β-gal within or frequency of ECs in mice with *Dnmt3a*^R878H/+^ vs. control hematopoiesis (Supplemental Figure 2G), further demonstrating that senescence occurs selectively within MSCs in the BM microenvironment. Increased MSC senescence was not observed upon transplant of *Dnmt3a*^R878H/+^ BM cells directly into non-conditioned aged mice (Supplemental Figure 2H-L). As SA-β-gal levels in MSCs were higher in old mice irrespective of being transplanted with control or *Dnmt3a*^R878H/+^ BM cells (Figure 2M and Supplemental Figure 2J), this suggests that substantive MSC senescence has already occurred as a natural consequence of aging and supports that *Dnmt3a*^R878H/+^ hematopoietic cells have the capacity to induce premature senescence in adult mice. Taken together, these data demonstrate that *Dnmt3a*^R878H/+^ hematopoietic cells induce MSC senescence *in vivo*, even with a relatively small proportion of mutant hematopoietic cells (equivalent to a VAF of 1-4%).

### Induction of BMSC Senescence by *Dnmt3a*-Mutant Hematopoietic Cells is Contact-Independent

To test the extent to which MSC senescence is caused directly by *Dnmt3a*^R878H/+^ HSPCs, we used an *ex vivo* co-culture system. Primary wild-type MSCs were isolated from mice and co-cultured with *Dnmt3a*^R878H/+^ or control HSPCs for 7 days (Figure 3A). MSCs co-cultured with *Dnmt3a*^R878H/+^ vs. control HSPCs had increased intracellular SA-β-gal (Figure 3B and Supplemental Figure 3A), and increased transcript expression of the senescence markers *Cdkn2a* (p16) and *Cdkn1a* (p21) (Figure 3C). In addition, MSCs co-cultured with *Dnmt3a*^R878H/+^ vs. control HSPCs had increased transcript expression of the SASP components *Il6, Tnf* and *Il1a* (Figure 3D). Analysis of the hematopoietic cell fraction after co-culture revealed an increase in mature myeloid cells and decrease in MPP^G/M^ and HSPCs in *Dnmt3a*^R878H/+^ compared to control cells (Figure 3E). Co-culture of mouse ECs with *Dnmt3a*^R878H/+^ or control HSPCs resulted in no change in SA-β-gal (Supplemental Figure 3B-D), further demonstrating the selectivity of senescence induction to MSCs. These data show that *Dnmt3a*^R878H/+^ HSPCs directly induce senescence of MSCs *ex vivo* and this is associated with increased myeloid cell differentiation.

**Figure 3.**
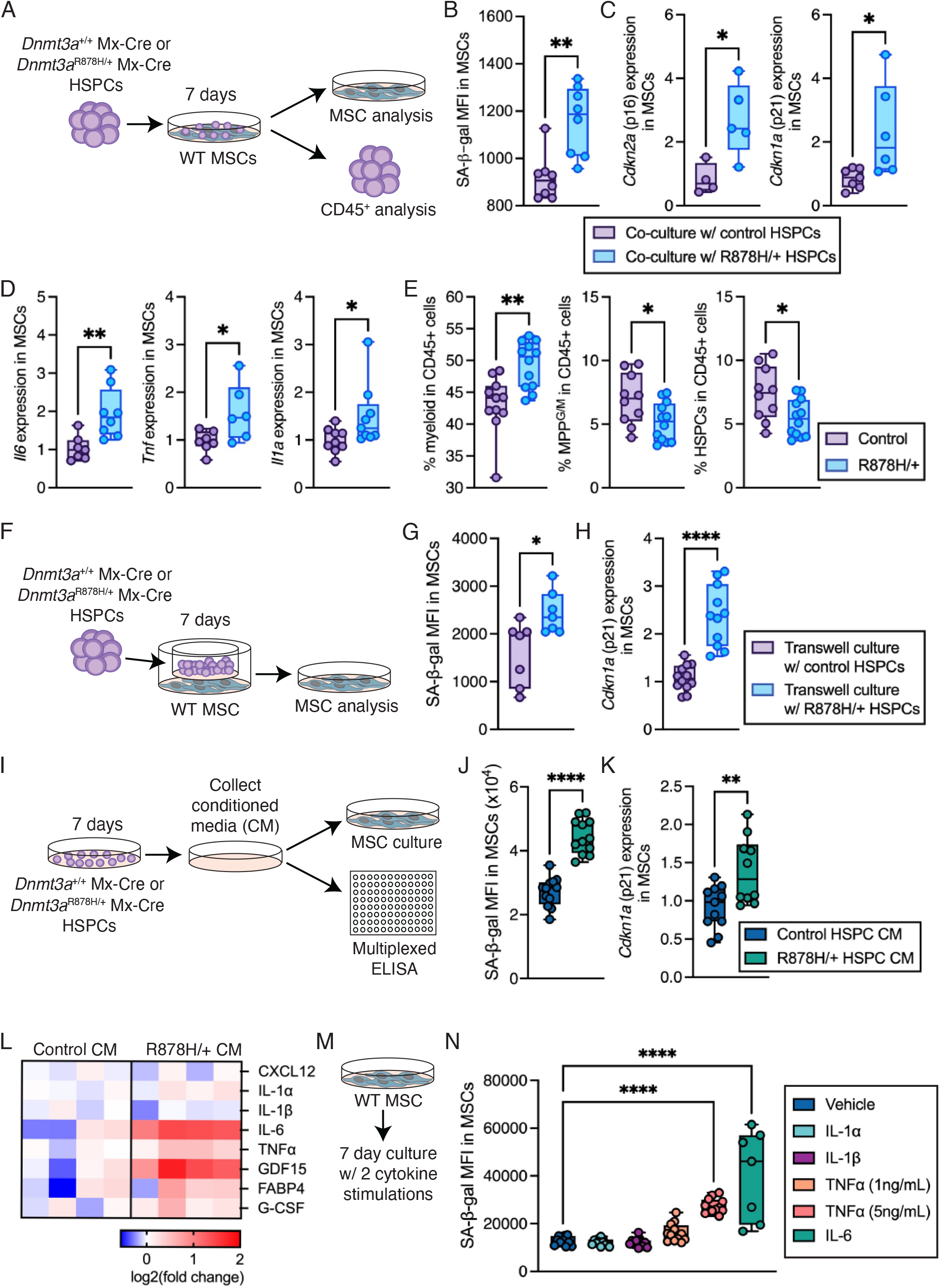
Soluble factors produced by *Dnmt3a*-mutant hematopoietic progenitor cells induce MSC senescence *ex vivo*. (**A**) Schematic of co-culture experimental design. (**B**) MFI of SA-β-gal in wild-type (WT) MSCs after co-culture with control or *Dnmt3a*^R878H/+^ HSPCs. \*\**P* < 0.01 by unpaired *t* test (*n* = 8). (**C**) Relative expression of *Cdkn2a* and *Cdkn1a* in MSCs after co-culture with control or *Dnmt3a*^R878H/+^ HSPCs. \**P* < 0.05 by Mann-Whitney test (*n* = 4-7). (**D**) Relative expression of *Il6, Tnf* and *Il1a* in MSCs after co-culture with control or *Dnmt3a*^R878H/+^ HSPCs. \**P* < 0.05, \*\**P* < 0.01 by Mann-Whitney test (*n* = 6-9). (**E**) Frequency of myeloid, MPP^G/M^ and HSPCs in CD45+ cells produced from control or *Dnmt3a*^R878H/+^ HSPCs after co-culture with WT MSCs. \**P* < 0.05, \*\**P* < 0.01 by unpaired *t* test (*n* = 10-12). (**F**) Schematic of transwell experimental design. (**G**) MFI of SA-β-gal in MSCs following transwell culture with control or *Dnmt3a*^R878H/+^ HSPCs. \**P* < 0.05 by Mann-Whitney test (*n* = 7). (**H**) Relative expression of *Cdkn1a* in MSCs following transwell culture with control or *Dnmt3a*^R878H/+^ HSPCs. \*\*\*\**P* < 0.0001 by Mann-Whitney test (*n* = 12). (**I**) Schematic of conditioned media (CM) experimental design. (**J**) MFI of SA-β-gal in wild-type (WT) MSCs after culture with CM from control or *Dnmt3a*^R878H/+^ HSPCs. \*\*\*\**P* < 0.0001 by unpaired *t* test (*n* = 12). (**K**) Relative expression of *Cdkn1a* in MSCs after culture with CM from control or *Dnmt3a*^R878H/+^ HSPCs. \*\**P* < 0.01 by unpaired *t* test (*n* = 10-11). (**L**) Log_2_FC in concentration of cytokines and growth factors in BM from control or *Dnmt3a*^R878H/+^ HSPCs assessed by multiplexed ELISA. *n* = 4. (**M**) Schematic of cytokine stimulation experimental design. (**N**) MFI of SA-β-gal in MSCs following culture with individual recombinant cytokines. \*\*\*\**P*<0.0001 by one-way ANOVA with Dunnett’s multiple comparisons test (*n* = 7-12).

To evaluate the cell contact-dependent vs.-independent induction of senescence of MSCs by *Dnmt3a*^R878H/+^ HSPCs, we established an *ex vivo* transwell co-culture assay (Figure 3F). Primary wild-type MSCs isolated from mice were co-cultured with *Dnmt3a*^R878H/+^ or control HSPCs in the transwell assay for 7 days. MSCs co-cultured with *Dnmt3a*^R878H/+^ vs. control HSPCs had increased intracellular SA-β-gal (Figure 3G), and increased transcript expression of *Cdkn1a* (Figure 3H). These data suggest that MSC senescence is induced by *Dnmt3a*^R878H/+^ HSPCs in a cell contact-independent manner. To evaluate this using an independent experimental strategy, we generated conditioned media (CM) by culturing *Dnmt3a*^R878H/+^ or control HSPCs alone for 7 days, then treated wild-type MSCs with this CM for 7 days (Figure 3I). MSCs cultured with *Dnmt3a*^R878H/+^ HSPC-CM had increased intracellular SA-β-gal (Figure 3J) and *Cdkn1a* expression (Figure 3K).

Given that *Dnmt3a*^R878H/+^ HSPCs induce MSC senescence in a cell contact-independent manner, we quantitated soluble factors produced by *Dnmt3a*^R878H/+^ HSPCs that may have the property of senescence induction. Using a Luminex assay, *Dnmt3a*^R878H/+^ HSPC-CM contained elevated levels of IL-6, TNFα, GDF-15, G-CSF and FABP4 but not CXCL12, IL-1α, IL-1β or G-CSF compared to control HSPC-CM (Figure 3L). A subset of these soluble factors (IL-1α, IL-1β, IL-6 and TNFα) were evaluated for their ability to induce senescence in primary wild-type MSCs (Figure 3M). MSCs treated with recombinant IL-6 or TNFα for 7 days had increased intracellular SA-β-gal compared to vehicle control treatment (Figure 3N). Together, these data show that *Dnmt3a*^R878H/+^ HSPCs produce soluble factors including IL-6 and TNFα which induce MSC senescence in a cell contact-independent manner.

### Depletion of Senescent Cells Using Navitoclax Reduces *Dnmt3a-*Mutant Hematopoiesis

Next, we tested the extent to which targeted removal of senescent cells would be sufficient to reduce *Dnmt3a*^R878H/+^ hematopoiesis. Given that MSC senescence induced by *Dnmt3a*^R878H/+^ hematopoiesis is characterized by increased levels of the anti-apoptotic proteins BCL-xL and BCL-2 (Figure 2G), we elected to utilize the senolytic navitoclax (ABT-263). Navitoclax targets BCL-xL, BCL-2 and BCL-W(*26*) and has been shown to deplete senescent HSCs following total body irradiation reversing aging of the hematopoietic system(*27*). We transplanted control or *Dnmt3a*^R878H/+^ BM into young wild-type CD45.1^+^ recipient mice following busulfan conditioning (Figure 4A). At 12 weeks post-transplant, we randomly distributed mice into vehicle control or navitoclax treatment groups (Supplemental Figure 4A) and followed an intermittent dosing schedule of once daily for 7 days followed by 14 days off, repeated twice (*27*) (Figure 4A). In mice with control hematopoiesis, navitoclax treatment did not alter SA-β-gal staining of MSCs (Figure 4B) or the proportion of BCL-xL+ or BCL-2+ MSCs (Supplemental Figure 4B). There was also no change in donor-derived BM hematopoietic cell engraftment (Figure 4C), or the proportion of myeloid cells (Figure 4D), HSPCs (Figure 4E), MPP^G/M^ (Figure 4F) or HSCs (Figure 4G) in donor-derived BM. In contrast, in mice with *Dnmt3a*^R878H/+^ hematopoiesis, navitoclax treatment resulted in reduced SA-β-gal staining of MSCs (Figure 4H) and the proportion of BCL-xL+ MSCs (Supplemental Figure 4C). We also observed reduced donor-derived BM hematopoietic cell engraftment (Figure 4I), decreased proportion of myeloid cells (Figure 4J), HSPCs (Figure 4K) and MPP^G/M^ (Figure 4L) in donor-derived BM. Short-term intermittent navitoclax treatment did not alter the proportion of HSCs (Figure 4M). To independently assess the proportion of myeloid progenitor cells, we performed myeloid colony-forming unit (CFU) assays using whole BM from *Dnmt3a*^R878H/+^ or control transplant mice. In mice with control hematopoiesis, navitoclax treatment did not change primary (passage 1) or secondary (passage 2) CFU formation (Figure 4N). In contrast, navitoclax treatment of mice with *Dnmt3a*^R878H/+^ hematopoiesis resulted in reduced primary and secondary CFU formation compared to vehicle treatment (Figure 4N). Together, removal of senescent cells *in vivo* reduces *Dnmt3a*^R878H/+^ hematopoiesis, particularly with respect to myeloid lineage cells.

**Figure 4.**
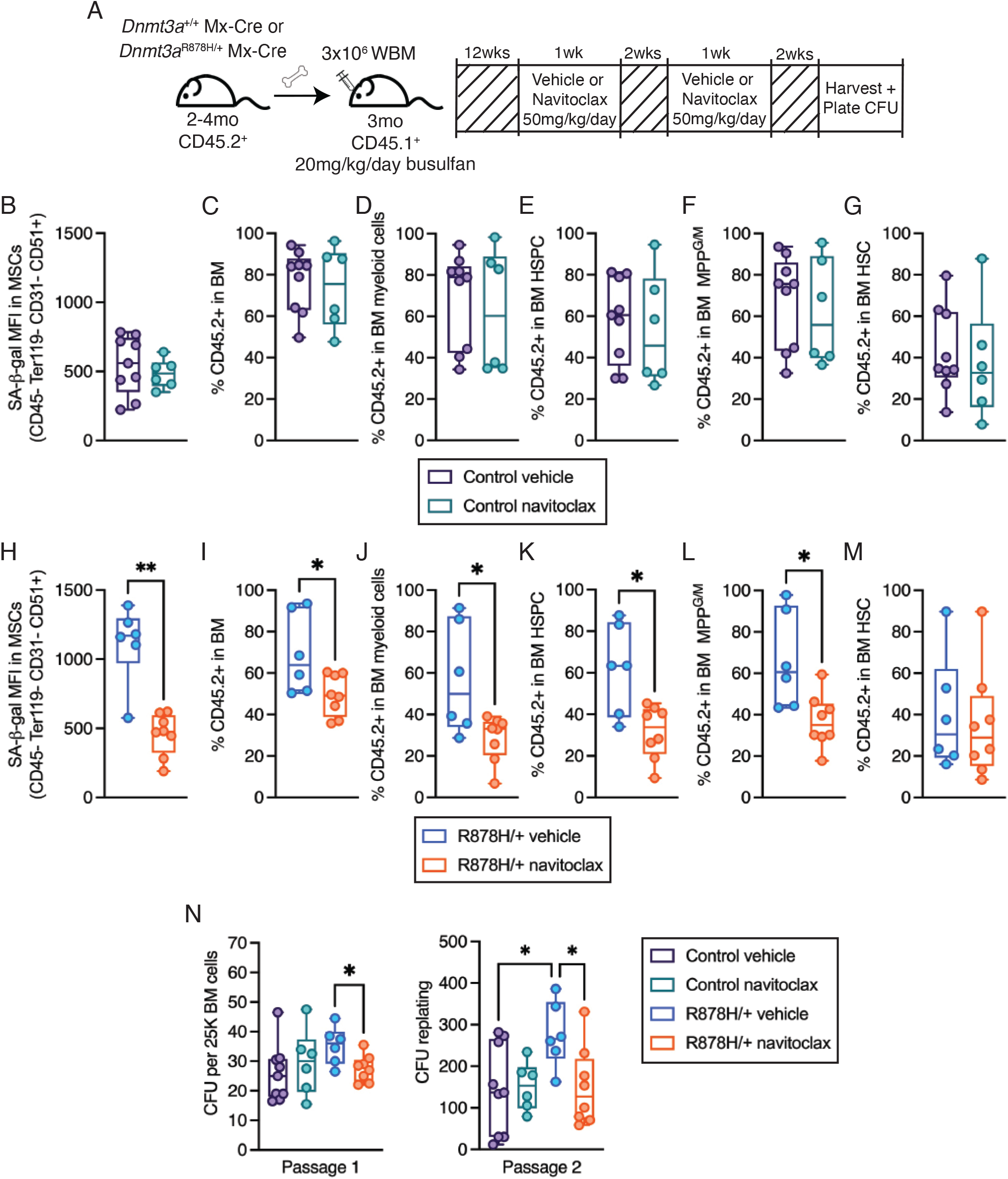
The senolytic navitoclax depletes senescent MSCs and reduces *Dnmt3a-*mutant hematopoiesis. (**A**) Schematic of navitoclax experimental design. (**B**) MFI of SA-β-gal in MSCs isolated from mice with control hematopoiesis after treatment with vehicle or navitoclax. (*n* = 6-9). (**C-G**) Frequency of donor (CD45.2^+^) cells in (**C**) BM, (**D**) myeloid cells, (**E**) HSPCs, (**F**) myeloid-primed multipotent progenitors (MPP^G/M^) and (**G**) HSCs in mice with control hematopoiesis after treatment with vehicle or navitoclax. *n* = 6-9. (**H**) MFI of SA-β-gal in MSCs isolated from mice with *Dnmt3a*^R878H/+^ hematopoiesis after treatment with vehicle or navitoclax (*n* = 6-9). \*\**P*<0.01 by Mann-Whitney test (*n* = 6-9). (**I-M**) Frequency of donor (CD45.2^+^) cells in (**I**) BM, (**J**) myeloid cells, (**K**) HSPCs, (**L**) MPP^G/M^ and (**M**) HSCs in mice with *Dnmt3a*^R878H/+^ hematopoiesis after treatment with vehicle or navitoclax. \**P*<0.05 by Mann-Whitney test (*n* = 6-9). (**N**) Total colony forming units (CFU) at passage 1 and passage 2 from BM cells isolated from control or *Dnmt3a*^R878H/+^ mice after treatment with vehicle or navitoclax. \**P* < 0.01 by one-way ANOVA with Fisher’s LSD (*n* = 6-9).

Given that navitoclax was administered systemically, the experiments described above do not distinguish the effects of navitoclax on non-hematopoietic cells from the effects of navitoclax on *Dnmt3a*^R878H/+^ hematopoietic cells. Evaluating BCL-xL and BCL-2 protein levels in HSPCs and mature myeloid cells by flow cytometry demonstrates that *Dnmt3a*^R878H/+^ HSPCs and mature myeloid cells have similar proportions of BCL-xL+ and BCL-2+ cells compared to control HSPCs and mature myeloid cells (Supplemental Figure 4D). This data suggests that *Dnmt3a*^R878H/+^ hematopoietic cells should not be differentially sensitive to navitoclax. Supporting this, direct addition of navitoclax to myeloid CFU assays (Supplemental Figure 4E) demonstrates that navitoclax does not alter primary (passage 1), secondary (passage 2), or tertiary (passage 3) CFU formation from *Dnmt3a*^R878H/+^ or control HSPCs (Supplemental Figure 4F). Together, this shows that depletion of senescent, non-hematopoietic cells in mice with *Dnmt3a*^R878H/+^ hematopoiesis reduces the proportion of *Dnmt3a*^R878H/+^ hematopoietic cells, particularly impacting myeloid-primed multipotent progenitor cells and downstream myeloid lineage cells.

### Depletion of Senescent Cells Delays Progression to Myeloid Neoplasms

Mitigating the risk of progression from CH to hematologic malignancy is of unmet clinical need. Thus, we evaluated the extent to which targeted removal of senescent MSCs would reduce the progression of *Dnmt3a*^R878H/+^ hematopoiesis to myeloproliferative neoplasms. Our pre-clinical experimental paradigm aimed to evaluate treatment with the senolytic navitoclax at the CH stage followed by withdrawal of senolytic treatment and evaluation of the development of myeloid neoplasms (Figure 5A). This design models a placebo-controlled trial in which individuals with CH are administered short-term treatment with navitoclax. As mice have not been reported to spontaneously develop *Npm1*^cA^ mutations driving myeloid neoplasms, this study employed our sequentially inducible genetically engineered *Dnmt3a*^R878H/+^ *Npm1*^cA/+^ mouse model(*28, 29*). Briefly, cre recombinase expression was activated to induce *Dnmt3a*^R878H/+^, 12 weeks later the mice were randomly assigned to senolytic (navitoclax) or placebo (vehicle) treatment groups, and following treatment, the *Npm1*^cA/+^ mutation was induced in all mice using tamoxifen to drive disease progression(*28*).

**Figure 5.**
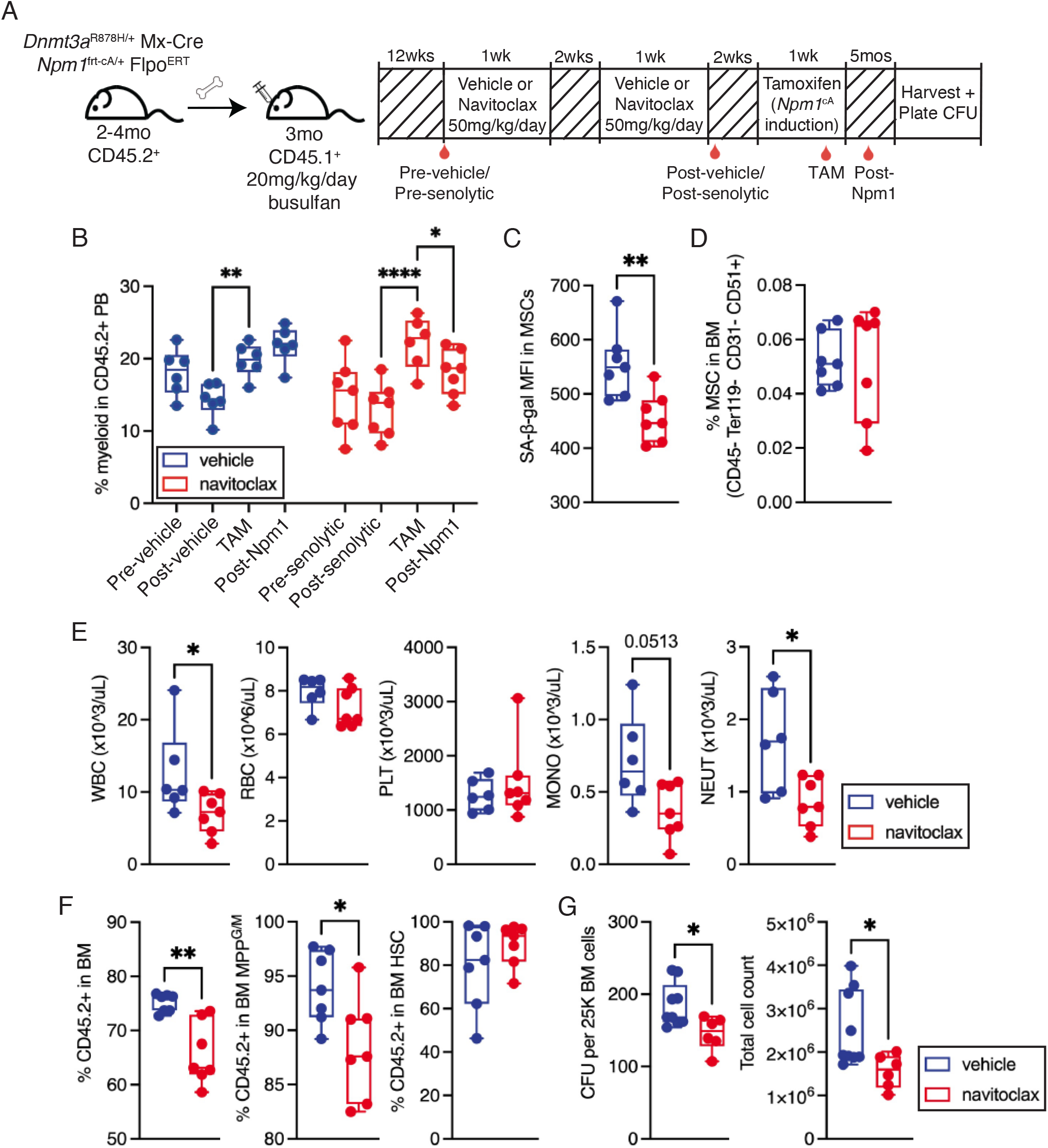
Clearance of *Dnmt3a*-mutant HSPC-induced MSC senescence reduces myeloproliferation of *Dnmt3a;Npm1*-mutant cells. (**A**) Schematic of experimental design evaluating the effect of navitoclax on progression from *Dnmt3a*^R878H/+^ clonal hematopoiesis to *Dnmt3a;Npm1*-mutant myeloproliferation. (**B**) Frequency of myeloid cells in donor-derived PB. \**P*<0.05, \*\**P*<0.01, \*\*\*\**P*<0.0001 by two-way ANOVA with Fisher’s LSD (*n =* 6-7). (**C**) MFI of SA-b-gal in MSCs isolated from mice with *Dnmt3a;Npm1*-mutant hematopoiesis treated with vehicle or navitoclax. \*\**P*<0.01 by Mann-Whitney test (*n* = 7). (**D**) Frequency of MSCs in the BM of mice with *Dnmt3a;Npm1*-mutant hematopoiesis treated with vehicle or navitoclax (*n* = 7). (**E**) Complete blood count values in mice with *Dnmt3a;Npm1*-mutant hematopoiesis treated with vehicle or navitoclax. WBC; white blood cells, RBC; red blood cells, PLT; platelets, MONO; monocytes, NEUT; neutrophils. \**P*<0.05 by Mann-Whitney test (*n* = 6-7). (**F**) Frequency of donor cells in BM, MPP^G/M^, and HSCs in mice with *Dnmt3a;Npm1*-mutant hematopoiesis treated with vehicle or navitoclax. \**P*<0.05, \*\**P*<0.01 by Mann-Whitney test (*n* = 7). (**G**) Total CFU and total cell count from CFU assay after passage 1 from BM cells isolated from *Dnmt3a;Npm1*-mutant mice treated with vehicle or navitoclax. \**P* < 0.01 by Mann-Whitney test (*n* = 6-9).

Examining the peripheral blood one week after induction of the *Npm1*^cA/+^ mutation using tamoxifen (“TAM”), mice in both the vehicle and navitoclax treatment groups had increased frequency of myeloid cells compared to baseline (Figure 5B). Four weeks later (“Post-Npm1”), vehicle-treated mice had similar levels of myeloid cells whereas navitoclax-treated mice had a reduced frequency of myeloid cells (Figure 5B). Harvesting mice 4 months later, the navitoclax-treated group had decreased SA-β-gal in BM MSCs (Figure 5C) compared to the vehicle-treated group without an observable change in the frequency of MSCs (Figure 5D). In addition, navitoclax-treated mice had significantly decreased WBC and neutrophil (NEUT) counts compared to vehicle-treated mice and trended toward decreased monocyte (MONO) count with no change in red blood cells (RBC) or platelets (PLT) (Figure 5E). In the BM, navitoclax-treated mice had a reduced proportion of *Dnmt3a;Npm1-*mutant hematopoietic cells and MPP^G/M^ cells compared to vehicle-treated mice, but no observed change in HSCs (Figure 5F). To assess proportions of functional myeloid progenitor cells, we performed myeloid CFU assays from donor-derived BM. The BM of navitoclax-treated *Dnmt3a;Npm1-*mutant mice generated fewer primary CFU and total myeloid cells compared to vehicle-treated mice (Figure 5G). While these mice did not develop a bona fide myeloid malignancy over the 5-month time course of the experiment, these results suggest that senolytic removal of senescent cells induced by *Dnmt3a*^R878H/+^ hematopoiesis can delay the progression of clonal hematopoiesis to myeloid neoplasms.

## Discussion

We demonstrate that *Dnmt3a*^R878H/+^ HSPCs directly alter the BM microenvironment and induce senescence selectively in MSCs, both in the context of steady state ‘native’ hematopoiesis as well as following transplantation. This is found even when the proportion of *Dnmt3a*^R878H/+^ HSPCs in the BM is small (equivalent to a human CH VAF of 1-4%), supporting the translational relevance of this observation. Senescence induction does not rely on physical cell-to-cell interaction but instead is caused by pro-inflammatory cytokines including IL-6 and TNFα which are produced by *Dnmt3a*^R878H/+^ HSPCs. Using the senolytic navitoclax, we show that MSC senescence contributes to the selective advantage of *Dnmt3a*^R878H/+^ HSPCs and their progeny. Further, depletion of senescent cells reduces myeloproliferation during the transformation of *Dnmt3a*^R878H/+^ HSPCs to myeloid neoplasms. These results establish new insights into how clones in CH can alter their surrounding BM microenvironment to create a favorable niche that facilitates their competitive advantage and promotes outgrowth of neoplastic clones.

Increased inflammation in the context of an aging or stressed BM microenvironment has been demonstrated in multiple studies to promote positive selection of CH-mutant HSCs(*20, 30, 31*). A complexity in understanding the mechanisms driving senescence induction by pro-inflammatory cytokines is that the cytokines produced by the ‘inducers’ of senescence (ie. *Dnmt3a*^R878H/+^ HSPCs) overlap with cytokines produced by senescent cells as part of the SASP. Thus, disentangling cause vs. consequence can be challenging. By integrating data across our experiments, we find that IL-6 and TNFα are produced by *Dnmt3a*^R878H/+^ HSPCs, individually are sufficient to induce MSC senescence, and are elevated at the protein level in the BM microenvironment of mice with *Dnmt3a*^R878H/+^ hematopoiesis. These data support that IL-6 and TNFα are a cause of MSC senescence but do not rule out the possibility that they are also released by senescent MSCs as part of the SASP. Increased TNFα signaling has been associated with CH in murine models(*20*) and in humans(*32*) as a key driver in *Dnmt3a*-mutant hematopoiesis. Elevated IL-6 has also been associated with CH as well as in the progression from MDS to AML. Data from 4,229 individuals with CH found that the *DNMT3A* mutation was associated with elevated plasma IL-6(*33*), and administration of anti-IL-6 neutralizing antibodies was found to reduce the selective growth advantage of *Dnmt3a*-mutant HSCs in fatty bone marrow(*34*). Increased levels of the IL-6 receptor were found in patients with high-risk MDS and antibody-based blocking of IL-6 signaling in a mouse model of MDS prevented AML progression(*35*). Interestingly, IL-6 is also associated with other CH-related pathologies such as cardiovascular disease. Heritable genetic deficiency in IL-6 production reduced the risk of cardiovascular disease in individuals with CH(*36*) and IL-6 receptor antibody treatment reversed atherosclerosis in a mouse model of CH driven by mutation in *Tet2*(*37*). The extent to which circulating CH-mutant hematopoietic cells induce senescence in other tissues, and contribute to CH-associated disease states, will be an exciting area of further investigation.

Involvement of the BM microenvironment in CH pathogenesis builds upon robust literature demonstrating that alterations in the microenvironment drive hematologic malignancies. MDS cells have been shown to reprogram the molecular characteristics of MSCs, creating an inflammatory milieu to which they are resistant and a favorable environment for their preferential growth(*38*). Deletion of *Dicer1* selectively in mesenchymal osteoprogenitors can induce MDS and AML(*39*), and an activating mutation in β-catenin in osteoblasts leads to development of AML(*40*). Deficiency of retinoic acid receptor gamma (RARγ)(*41*) or retinoblastoma (RB)(*42*) in the BM microenvironment causes development of myeloproliferative disorder. A case study report of multiple donor-derived leukemias in a recipient of multiple allogeneic BM transplants supports the concept that changes in the BM microenvironment can create a niche for leukemogenesis and that a healthy BM microenvironment can restrain a genetically abnormal HSC(*43*). Extending this concept to pre-malignant states, we propose that priming of the microenvironment by CH-mutant cells can promote clonal evolution to hematologic malignancies.

Navitoclax (ABT-263) has been shown to directly target aged HSCs and progenitor cells(*27*). We find that *Dnmt3a*^R878H/+^ HSCs and progenitor cells in young mice are not directly depleted by navitoclax and do not express markers of senescence. Thus, *Dnmt3a*^R878H/+^ HSCs do not undergo ‘accelerated aging’ with respect to these markers, and navitoclax reduces *Dnmt3a*-mutant hematopoiesis and progression to myeloid neoplasms independent of direct effects on the hematopoietic system. Given that navitoclax is administered to mice systemically, our experimental data does not rule out that navitoclax may impact other non-hematopoietic cell populations apart from MSCs. Broadly, senolytics have shown promise in the treatment of age-related disease(*44*).

New senolytics continue to be identified using high-throughput library screens and other approaches(*45*), and intermittent administration appears as effective as continuous treatment for depleting senescent cells(*46*). This intermittent treatment strategy may help in reducing adverse effects in the case of compounds like navitoclax which causes dose-dependent thrombocytopenia(*47*). Given that senescent cells naturally accumulate during aging, senolytics could theoretically achieve anti-CH and anti-aging interventions(*48*). In preclinical models, interventions targeting senescent cells have been shown to delay, prevent or alleviate multiple disorders including diabetes, cardiac dysfunction, cognitive impairment, and osteoporosis(*44*).

Our findings in CH extend to other tissue types which accumulate somatic mutations in the context of aging. For example, in tissues such as the colon, endometrium, esophagus and skin, clonal evolution can occur years or decades prior to development of disease(*49*). Investigating how the stroma dynamically co-evolves with age in mutated but histologically normal clonal tissues, including induction of senescence, is of increasing interest. Emerging efforts such as “The PreCancer Atlas (PCA)” aim to systematically analyze longitudinal clonal architecture of premalignant lesions within all major tissues at cell-intrinsic (genomic, transcriptomic, and epigenetic) and cell-extrinsic (stromal and immune) levels(*50, 51*). Through characterization of the synergies between molecular alterations in premalignant lesions and changes in the microenvironment associated with progression, the goal is to develop biomarkers for early detection and risk stratification as well as identify preventive interventions to reverse or delay the development of cancer. While great effort has focused on changes in the immune cell microenvironment and repression of functional immune cells by activation of immune checkpoints(*52*), our work supports that the non-hematopoietic stromal cell niche is also a major driver of clonal evolution and transformation. Thus, targeting the stromal cell niche represents a novel therapeutic strategy to delay or prevent transformation from CH to hematologic malignancy.

## Methods

### Mice

C57BL/6J (The Jackson Laboratory (JAX) stock #00664, referred to as CD45.2^+^) and B6.SJL-*Ptprc^a^ Pepc^b^*/BoyJ (JAX stock #002014, referred to as CD45.1^+^) mice were obtained from and aged within JAX. *Dnmt3a*^fl-R878H/+^ mice(*28*) (JAX stock #032289) were crossed to B6.Cg-Tg(Mx1-cre)1Cgn/J mice(*53*) (JAX stock #003556, referred to as Mx-Cre) or C57BL/6N*-Fgd5^tm3(cre/ERT2)Djr^*/J mice(*54*) (JAX stock #027789, referred to as Fgd5-Cre). Mice carrying the Mx-Cre allele were given 15Lmg/kg polyinosinic-polycytidylic acid (poly(I:C)) (InvivoGen) intraperitoneally every other day for a total of five doses between 2-4 months of age, prior to use. Mice carrying the Fgd5-Cre allele were given 125Lmg/kg tamoxifen by oral gavage every day for a total of 3 doses between 2-4 months of age, prior to use. Mice were housed under specific pathogen-free conditions in a 12/12-hour light/dark cycle with food and water provided *ad libitum*. Mice used were at 2-4 months of age (young) or 12 months of age (aged) and both biological sexes were used for experiments except for transplantation experiments where female mice were used. All animal work used in this study was approved by The Jackson Laboratory’s Institutional Animal Care and Use Committee.

### Bone marrow transplantation

For transplantation experiments into middle-aged CD45.1^+^ recipient mice, 3x10^6^ whole bone marrow from *Dnmt3a*^+/+^ Fgd5-CreER^T2^ LSL-tdTomato or *Dnmt3a*^R878H/+^ Fgd5-CreER^T2^ LSL-tdTomato donor mice were transplanted retro-orbitally into 9-month-old busulfan-conditioned mice (20mg/kg/per day interperitoneally for consecutive 3 days). For transplantation experiments into young CD45.1^+^ recipient mice, 3x10^6^ whole bone marrow from *Dnmt3a*^+/+^ Mx-Cre or *Dnmt3a*^R878H/+^ Mx-Cre donor mice were transplanted retro-orbitally into 3-month-old busulfan-conditioned mice. For non-conditioned transplant experiments, over 3 consecutive days, 5x10^6^ whole bone marrow cells from *Dnmt3a*^+/+^ Mx-Cre or *Dnmt3a*^R878H/+^ Mx-Cre were injected into non-conditioned young (2 months old) or aged (13 months old) CD45.1^+^ recipients via retro-orbital injection. For transplantation experiments assessing AML initiation, 3x10^6^ whole bone marrow from *Dnmt3a*^R878H/+^ *Npm1*^frt-cA/+^ Mx-Cre *R26*^FlpoER^ were transplanted retro-orbitally into 3-month-old busulfan-conditioned mice. Donor mice received poly(I:C) prior to transplant. 12 weeks post-transplantation, mice were treated with navitoclax or vehicle control. Two weeks later, mice received 125Lmg/kg tamoxifen by oral gavage every day for a total of 3 doses to induce *Npm1*^cA/+^ mutation. Navitoclax (in ethanol:polyethylene glycol 400:Phosal 50 PG at a ratio of 10:30:60) was administered to mice by oral gavage at 50 mg/kg/per day once daily for 7 days followed by 14 days off, repeated twice.

### Bone marrow isolation and cell sorting for single cell RNA-sequencing

For hematopoietic cell isolation, single-cell suspensions of BM were prepared by filtering crushed femurs, tibias, and iliac crests from each mouse. Red cells were lysed using 1X red blood cell lysis buffer (ThermoFisher). Cells were then stained with a combination of fluorochrome-conjugated antibodies from eBioscience, BD Biosciences, BioLegend or Miltenyi: CD45.1 (clone A20), CD45.2 (clone 104), c-Kit (clone 2B8), Sca-1 (clone 108129), CD150 (clone TC15-12F12.2), CD48 (clone HM48-1), FLT3 (clone A2F10), CD34 (clone RAM34), FcγR (clone 2.4G2), a mature lineage (Lin) marker mix including B220 (clone RA3-6B2), CD11b (clone M1/70), CD4 (clone RM4-5), CD5 (clone 53-7.3), CD8a (clone 53-6.7), Ter-119 (clone TER-119) and Gr-1 (clone RB6-8C5), and a viability stain.

For isolation of non-hematopoietic cells, femurs, tibias, and iliac crests were cut in half then spun into 100 uL of PBS. The cell pellet was resuspended in 5mL digest solution (2mg/mL collagenase IV and 1mg/mL dispase in 1X HBSS) in one well of 6 well plate. Bone chips were placed into 5mL digest solution in one well of 6 well plate. The plate was incubated at 37°C in 5% CO_2_ for 10 minutes. Bone chips were mixed, digest solution was removed from the bone marrow plug and added to ice cold FACS buffer and replaced with fresh 5mL of digest solution. This was repeated 3 times and all digest buffer was pooled into ice cold FACS buffer and filtered. Cells were then stained with a combination of fluorochrome-conjugated antibodies from eBioscience, BD Biosciences, BioLegend or Miltenyi: CD45 (clone 30-F11), Ter-119 (clone TER-119), CD41 (clone MWReg 30), CD31 (clone MEC 13.3), CD71 (clone R17217), CD51 (clone RMV-7), and a viability stain.

From each individual mouse, hematopoietic and non-hematopoietic cell fractions were kept separate and serially sorted into the same collection tube. First, hematopoietic cell populations were sorted on a FACSAria II (BD): 5000 mature hematopoietic cells (CD45.2^+^ tdTomato^+^ Lin^+^), 6000 committed hematopoietic progenitors (CD45.2^+^ tdTomato^+^ Lin^-^ c-Kit^+^ Sca1^-^), and 7000 hematopoietic stem and multipotent progenitor cells (CD45.2^+^ tdTomato^+^ Lin^-^ c-Kit^+^ Sca1^+^) into a 1.5ml tube with 10ul IMDM + 10% FBS at 4°C. Next, this tube was transferred to sort non-hematopoietic cell populations using a FACSymphony S6 (BD): 2000 endothelial cells (CD45.2^-^ Ter119^-^ CD41^-^ CD31^+^) and 8000 mesenchymal stromal cells (CD45.2^-^ Ter119^-^ CD41^-^ CD31^-^ CD71^-^ CD51^+^). Cells were immediately counted on a Countess II automated cell counter (ThermoFisher) and 12,000 cells were loaded on to one lane of a 10X Chromium microfluidic chip (10X Genomics).

### Single cell RNA-sequencing

Single cell capture, barcoding, and library preparation was performed using 10X Chromium version 3.1 NEXTGEM chemistry according to the manufacturer’s protocol (#CG000315). Prior to sequencing, quality control of the cDNA libraries was assessed using an Agilent 4200 Tapestation and quantified by KAPA qPCR. Libraries were sequenced using an Illumina NovaSeq 6000 S4 flow cell lane targeting 6,000 barcoded cells with an average sequencing depth of 75,000 reads per cell. Illumina base call files for all libraries were demultiplexed and converted to FASTQs using bcl2fastq v2.20.0.422. The Cell Ranger pipeline (10X Genomics) was used to align reads to the mouse reference GRCm38.p93 (mm10 10X Genomics reference 2020-A), deduplicate reads, call cells, and generate cell by gene digital count matrices for each library. The matrices were uploaded into PartekFlow (version 10.0.23.0720) for downstream analysis and visualization including log transformation of count data, principal component analysis, and graph-based clustering from the top 18 principal components. This was using the Louvain algorithm, t-distributed stochastic neighbor embedding (t-SNE) visualization, and pathway enrichment analysis. The 30 top marker genes in each of the clusters were cross-referenced with published single cell RNA-seq profiles to annotate each population.

### Flow cytometry cell surface marker profiles

The following surface marker combinations were used for cell sorting and/or analysis: HSC (Lin^-^ Sca-1^+^ c-Kit^+^ Flt3^-^ CD150^+^ CD48^-^), MPP^G/M^ (Lin^-^ Sca-1^+^ c-Kit^+^ Flt3^-^ CD150^-^ CD48^+^), LSK (Lin^-^ Sca-1^+^ cKit^+^), MyPro (Lin^-^ Sca-1^-^ c-Kit^+^), B cells (B220^+^ CD11b^-^ CD3e^-^), T cells (CD3e^+^ B220^-^ CD11b^-^), myeloid cells (CD11b^+^ B220^-^ CD3^-^), granulocytes (CD11b^+^ B220^-^ CD3^-^ Ly6g^+^ Ly6c^+^), MSCs (CD45^-^ CD41^-^ Ter119^-^ CD31^-^ CD51^+^) and endothelial cells (CD45^-^ CD41^-^ Ter119^-^ CD31^+^). Antibody cocktails were incubated with BM cells for >30 min at 4°C. When using the β-galactosidase stain C12FDG (5-Dodecanoylaminofluorescein Di-β-D-Galactopyranoside, ThermoFisher), cells were incubated with C12FDG dye for 2 hoursLat 37°C and washed twice before staining with surface marker antibodies. For experiments using BCL-xL (clone 54H6) or BCL-2 (clone REA356), BM cells were fixed and permeabilized using the FIX & PERM™ Cell Permeabilization Kit (ThermoFisher) per the manufacturer’s instructions. Stained cells were sorted using a FACSAria II or a FACSymphony S6 Sorter or analyzed on a FACSymphony A5. FlowJo V10 was used for data analysis.

### Peripheral blood analysis

Peripheral blood was collected from mice via retro-orbital bleed. Red blood cells were lysed, and cells were stained with CD45.1 (clone A20), CD45.2 (clone 104), B220 (clone RA3-6B2), CD3e (clone 145-2C11), CD11b (clone M1/70), Ly6g (clone 1A8), Ly6c (clone HK1.4), and Gr-1 (clone RB6-8C5). Data was captured using a LSRII (BD) and analyzed using FlowJo V10. Complete blood counts were performed on whole blood using an Advia 120 Hematology Analyzer (Siemens).

### Cell culture

Cell culture was carried out in 5% CO_2_ in a humidified incubator at 37°C. MSCs were isolated from the mouse bone marrow by adherence to tissue culture plastic and expanded in minimum essential media (MEM) containing 20% FBS plus 1% penicillin-streptomycin as described (*55*). MSCs were evaluated by morphology and flow cytometry for confirmation of the phenotypic markers CD51^+^ CD31^-^ Ter119^-^ CD45^-^. Hematopoietic stem and progenitor cells (HSPCs) were isolated by positive selection using CD117 MicroBeads (Miltenyi cat. #130-091-224). Endothelial cells (ECs) were a kind gift from Dr. Jason Butler, University of Florida. ECs were cultured in a 1:1 mixture of low glucose DMEM and Ham’s F-12 (CellGro) supplemented with 20% heat-inactivated FBS, antibiotic-antimycotic, non-essential amino acids, 20 mM HEPES, 100 μg/mL heparin, 50 μg/mL endothelial growth factor and 1% Pen/Strep. MSCs or ECs were seeded at a density of 75,000 cells in a 24-well plate in normal growth media. Once 70% confluent, 0.1 × 10^6^ HSPCs were co-cultured with MSCs or ECs for 7 days. HSPCs were removed and analyzed by flow cytometry. MSCs were isolated using trypsin and analyzed by flow cytometry or RNA was isolated for qPCR. For transwell assays, MSCs were seeded at a density of 75,000 cells in a 24-well plate in normal growth media. Once 70% confluent, 0.1 × 10^6^ HSPCs were cultured in transwells on top of the MSCs for 7 days. MSCs were then analyzed by flow cytometry or RNA was extracted for qPCR. For conditioned media, HSPCs were isolated and cultured in StemSpan SFEM II (StemCell Technologies cat. #09605) with 10 ng/mL recombinant murine SCF (STEMCELL Technologies; cat. #78064) for 7 days. Cells were centrifuged at 1,500 rpm for 10 min at 4°C to isolate conditioned media. Conditioned media was added to 75,000 MSCs in a 24-well plate for 7 days and MSCs were then analyzed by flow cytometry and qPCR. For cytokine treatment, MSCs were seeded at a density of 75,000 cells in a 24-well plate in normal growth media. MSCs were treated with 1 ng/mL or 5 ng/mL recombinant murine TNFα (PeproTech; cat. #315-01A), 50 ng/mL recombinant IL-6 (StemCell Technologies cat. #78052.1) 3 ng/mL recombinant murine IL-1α (PeproTech; cat. #211-11A), or 3 ng/mL recombinant murine IL-1β (PeproTech; cat. #211-11B) for 7 days and analyzed by flow cytometry.

### Colony forming unit assay

Whole bone marrow from transplanted mice or HSPCs from *Dnmt3a*^+/+^ or *Dnmt3a*^R878H/+^ mice were plated in MethoCult GF M3434 (StemCell Technologies cat. #03434) at the indicated numbers and cultured at 37°C and 5% CO_2_. Colonies were scored 7-10 days post-plating using a Nikon Eclipse TS100 inverted microscope. For the CFU replating assay, colonies were harvested and 15Lx10^4^ cells were replated in fresh MethoCult GF M3434.

### Luminex analysis

One femur and one tibia from each mouse were cut in half, then spun into 100μL of PBS. For experiments using conditioned media, bone marrow isolation was prepared by isolating, filtering and crushed pooled tibias, femurs, and iliac crests of each mouse. HSPCs were isolated using CD117 MicroBeads (Miltenyi cat. #130-091-224) and cultured in StemSpan SFEM II (StemCell Technologies cat. #09605) with 10 ng/mL recombinant murine SCF (STEMCELL Technologies; cat. #78064) for 7 days. Cells were centrifuged at 1,500 rpm for 10 min at 4°C to isolate conditioned media. Custom Luminex Assays were used to assess analyte levels including mouse GDF15, IL-6, TNFα, IL-1α, IL-1β, MMP3, Osteopontin, G-CSF and CXCL12 (R&D Systems).

### Real-time PCR for senescence genes

RNeasy Micro Kit (Qiagen; cat. #74004) was used to extract whole cell RNA. qRT-PCR was performed using Power SYBR™ Green PCR Master Mix (Applied Biosystems; cat. #4368577). PCR conditions were pre-amplification (50 °C/120Lseconds, 95 °C/10 minutes), amplification for 40 cycles (95 °C/15Lseconds, 60 °C/60Lseconds), and melt curve (95 °C/15Lseconds, 60 °C/60Lseconds 95L°C/15Lseconds) on a QuantStudio 7 Flex (ThermoFisher). Messenger RNA (mRNA) expression was normalized against Hypoxanthine Guanine Phosphoribosyltransferase (HPRT) using the comparative cycle threshold method. Custom primers designed and purchased from Integrated DNA Technologies Inc. Sequences are shown in Supplemental Table 2.

### Quantification and statistical analysis

No group randomization or blinding was performed, and no statistical methods were used to determine sample size. All statistical tests were performed using Prism 9 software (GraphPad, RRID:SCR_002798). Due to variability in the data, statistical comparison of *in vivo* data with 2 groups was performed without assumption of normal distribution using Mann–Whitney test. For statistical comparison of more than two groups, 2way ANOVA or Kruskal–Wallis test followed by Dunn’s multiple comparisons or 2way analysis of variance was used. Differences among group means were considered significant when the probability value, *p*, was <0.05*, 0.01**, 0.001***. Sample size (*n*) represents number of biological replicates.

## Data availability

Raw single cell RNA-seq data is available through the Gene Expression Omnibus (GEO) accession GSE240686.

## Authorship

J.J.M. conceptualized the project. J.J.M. and J.J.T. designed experiments. J.J.M., K.A.Y, and P.A.C.D. performed experiments, and J.J.M., and J.J.T. analyzed data. I.F.M. and R.L.L. provided input on data interpretation. J.J.M. and J.J.T. wrote the manuscript. All authors edited the manuscript.

## Acknowledgements

This work was supported by National Institutes of Health grant U01AG077925 to J.J.T and R.L.L., and R01DK118072, R01AG069010, and an EvansMDS Discovery Research Grant to J.J.T. This work was supported in part by the NIH/NCI Cancer Center Support Grants P30CA034196 and P30CA008748. J.J.T. was supported by a Leukemia & Lymphoma Society Scholar Award and The Dattels Family Endowed Chair. J.J.M. is supported by a Leukemia & Lymphoma Society Career Development Program Fellow Award and The Jackson Laboratory Scholar Award. I.F.M. is supported by F99CA284253. We thank all members of the Trowbridge Lab for experimental support. We thank the The Jackson Laboratory’s Single Cell Biology, Genome Technologies, and Flow Cytometry Scientific Services. We thank Jason Butler for providing us with bone marrow endothelial cells. We thank Maria Telpoukhovskaia, Stuart Rushworth, Kristian Bowles, and Borhane Guezguez for their input into this work.

## SUPPLEMENTAL DATA

**Supplemental Figure 1. Analysis of single cell RNA-seq BM data from mice with control or *Dnmt3a*-mutant hematopoiesis.** (**A**) Frequency of 21 annotated hematopoietic cell clusters in mice with control or *Dnmt3a*^R878H/+^ hematopoiesis (*n* = 4). HSC/MPP: hematopoietic stem and multipotent progenitor cells, MPP^G/M^: granulocyte-macrophage-primed multipotent progenitor cells, MPP^Ly^: lymphoid-primed multipotent progenitor cells, MEP: megakaryocyte-erythroid progenitor cells, EryPro: erythroid progenitor cells, Ery: erythroblast, MEoBasoPro: mast-eosinophil-basophil progenitor cells, GMP: granulocyte-macrophage progenitor cells, NeutPro: neutrophil progenitor cells, ImmNeut: immature neutrophils, Neut: neutrophils, MonoPro: monocyte progenitor cells, Mono: monocytes, Mac: macrophages, Mono/DC: monocyte dendritic cell, DC: dendritic cells, PreB: pre-B cells, ProB: pro-B cells, B: B cells, NK/T: natural killer and T cells. (**B**) Gene set enrichment analysis of transcripts increased (red) and decreased (blue) in *Dnmt3a*^R878H/+^ versus control hematopoietic cell populations.

**Supplemental Figure 2. Additional parameters assessed in experiments interrogating MSC senescence induction by *Dnmt3a*-mutant hematopoiesis *in vivo***. (**A**) Representative flow cytometry gating of BM hematopoietic stem and progenitor cells. (**B**) Frequency of B, T, and myeloid cells in donor derived PB of mice with control or *Dnmt3a*^R878H/+^ hematopoiesis. Experimental design shown in Figure 2A. \**P*<0.05 by Mann-Whitney test (*n* = 8-9). (**C**) Frequency of granulocytes in donor-derived myeloid cells. \**P*<0.05 by Mann-Whitney test (*n* = 8-9). (**D**) MFI of SA-β-gal in endothelial cells (ECs) and frequency of ECs (*n* = 8-9). (**E**) Frequency of B, T, and myeloid cells in donor derived PB of mice at 60 weeks post non-conditioned transplant of control or *Dnmt3a*^R878H/+^ hematopoiesis. Experimental design shown in Figure 2I. \**P*<0.05 by Mann-Whitney test (*n* =7-8). (**F**) Frequency of granulocytes in donor-derived myeloid cells (*n* =7). (**G**) MFI of SA-β-gal in ECs and frequency of ECs (*n* =8). (**H**) Schematic of non-conditioned transplant design into aged recipient mice. (**I**) Frequency of donor cells in PB of aged recipient mice. \*\*\**P* < 0.001 by two-way ANOVA with Sidak’s multiple comparisons test (*n* = 8). (**J**) MFI of SA-β-gal in MSCs and frequency of MSCs in the BM of aged recipient mice at 60 weeks post-transplant (*n* =3-6). (**K**) MFI of SA-β-gal in ECs and frequency of ECs in the BM of aged recipient mice (*n* =3-6). (**L**) Frequency of donor cells in BM, HSPC and HSC in the BM of aged recipient mice (*n* = 3-6).

**Supplemental Figure 3. Lack of senescence induction of endothelial cells (ECs) by *Dnmt3a*-mutant hematopoietic progenitor cells *ex vivo*.** (**A**) Representative flow cytometry gating used to identify mesenchymal stromal cells (MSC) after co-culture with control or *Dnmt3a*^R878H/+^ HSPCs. (**B**) Representative flow cytometry gating used to identify endothelial cells (ECs) after co-culture with control or *Dnmt3a*^R878H/+^ HSPCs. (**C**) Schematic of co-culture experimental design. (**D**) MFI of SA-β-gal in wild-type (WT) ECs after co-culture with control or *Dnmt3a*^R878H/+^ HSPCs (*n* = 8-9).

**Supplemental Figure 4. The senolytic navitoclax does not directly target *Dnmt3a*-mutant hematopoietic stem and progenitor cells.** (**A**) Frequency of donor-derived cells in PB of mice with control or *Dnmt3a*^R878H/+^ hematopoiesis at 12 weeks post-transplant prior to administration of vehicle or navitoclax. Experimental design shown in Figure 4A. (*n* = 6-9). (**B**) Frequency of BCL-xL^+^ and BCL-2^+^ MSCs in the BM of mice with control hematopoiesis after treatment with vehicle or navitoclax. (*n* = 6-9). (**C**) Frequency of BCL-xL^+^ and BCL-2^+^ MSCs in the BM of mice with *Dnmt3a*^R878H/+^ hematopoiesis after treatment with vehicle or navitoclax. \**P*<0.05 by Mann-Whitney test (*n* = 6-8). (**D**) Frequency of BCL-xL^+^ and BCL-2^+^ cells in hematopoietic stem, progenitor, and mature cell compartments of control and *Dnmt3a*^R878H/+^ mice. *n* = 3. (**E**) Schematic of CFU experimental design testing the direct effect of navitoclax on *Dnmt3a*^R878H/+^ HSPCs. (**F**) Total CFU at passages 1, 2 and 3 control or *Dnmt3a*^R878H/+^ HSPCs treated directly with vehicle or navitoclax. \*\**P* < 0.01, \*\*\*\**P*<0.0001 by two-way ANOVA with Tukey’s multiple comparisons test (*n* = 3).

**Supplemental Table 1. Top 30 biomarkers defining hematopoietic and non-hematopoietic cell populations in single cell RNA-seq data.** See attached file.

**Supplemental Table 2.**
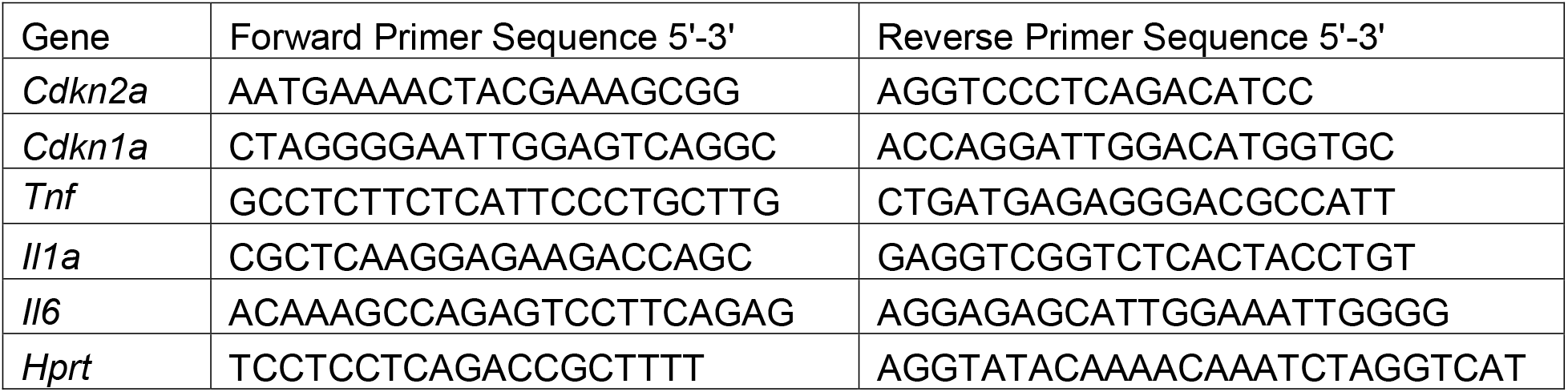
Primer sequences used for real-time PCR.

## REFERENCES

1. A. Mendelson, P. S. Frenette, Hematopoietic stem cell niche maintenance during homeostasis and regeneration. Nat Med 20, 833–846 (2014).

2. S. Jaiswal, B. L. Ebert, Clonal hematopoiesis in human aging and disease. Science 366, (2019).

3. S. Jaiswal et al., Age-related clonal hematopoiesis associated with adverse outcomes. N Engl J Med 371, 2488–2498 (2014).

4. L. D. Weeks, et al., Prediction of Risk for Myeloid Malignancy in Clonal Hematopoiesis. NEJM Evidence 2, (2023).

5. W. Fiedler et al., Vascular Endothelial Growth Factor, a Possible Paracrine Growth Factor in Human Acute Myeloid Leukemia. Blood 89, 1870–1875 (1997).

6. S. Dias et al., Inhibition of both paracrine and autocrine VEGF/ VEGFR-2 signaling pathways is essential to induce long-term remission of xenotransplanted human leukemias. Proc Natl Acad Sci U S A 98, 10857–10862 (2001).

7. S. Dias et al., Autocrine stimulation of VEGFR-2 activates human leukemic cell growth and migration. J Clin Invest 106, 511–521 (2000).

8. J. W. Hussong, G. M. Rodgers, P. J. Shami, Evidence of increased angiogenesis in patients with acute myeloid leukemia. Blood 95, 309–313 (2000).

9. M. Azadniv et al., Bone marrow mesenchymal stromal cells from acute myelogenous leukemia patients demonstrate adipogenic differentiation propensity with implications for leukemia cell support. Leukemia 34, 391–403 (2020).

10. H. Ye et al., Leukemic Stem Cells Evade Chemotherapy by Metabolic Adaptation to an Adipose Tissue Niche. Cell Stem Cell 19, 23–37 (2016).

11. M. S. Shafat et al., Leukemic blasts program bone marrow adipocytes to generate a protumoral microenvironment. Blood 129, 1320–1332 (2017).

12. E. D. Lagadinou et al., BCL-2 inhibition targets oxidative phosphorylation and selectively eradicates quiescent human leukemia stem cells. Cell Stem Cell 12, 329–341 (2013).

13. R. Moschoi et al., Protective mitochondrial transfer from bone marrow stromal cells to acute myeloid leukemic cells during chemotherapy. Blood 128, 253–264 (2016).

14. C. R. Marlein et al., NADPH oxidase-2 derived superoxide drives mitochondrial transfer from bone marrow stromal cells to leukemic blasts. Blood 130, 1649–1660 (2017).

15. Y. Zhao et al., Down-regulation of Dicer1 promotes cellular senescence and decreases the differentiation and stem cell-supporting capacities of mesenchymal stromal cells in patients with myelodysplastic syndrome. Haematologica 100, 194–204 (2015).

16. T. Andre et al., Evidences of early senescence in multiple myeloma bone marrow mesenchymal stromal cells. PLoS One 8, e59756 (2013).

17. P. Zhou et al., Senescent bone marrow microenvironment promotes Nras-mutant leukemia. J Mol Cell Biol 13, 72–74 (2021).

18. A. M. Abdul-Aziz et al., Acute myeloid leukemia induces protumoral p16INK4a-driven senescence in the bone marrow microenvironment. Blood 133, 446–456 (2019).

19. K. Young et al., Decline in IGF1 in the bone marrow microenvironment initiates hematopoietic stem cell aging. Cell Stem Cell 28, 1473–1482 e1477 (2021).

20. J. M. SanMiguel et al., Distinct Tumor Necrosis Factor Alpha Receptors Dictate Stem Cell Fitness versus Lineage Output in Dnmt3a-Mutant Clonal Hematopoiesis. Cancer Discov 12, 2763–2773 (2022).

21. C. Baccin et al., Combined single-cell and spatial transcriptomics reveal the molecular, cellular and spatial bone marrow niche organization. Nat Cell Biol 22, 38–48 (2020).

22. D. Hormaechea-Agulla et al., Chronic infection drives Dnmt3a-loss-of-function clonal hematopoiesis via IFNgamma signaling. Cell Stem Cell 28, 1428–1442 e1426 (2021).

23. C. R. Zhang et al., Txnip Enhances Fitness of Dnmt3a-Mutant Hematopoietic Stem Cells via p21. Blood Cancer Discov 3, 220–239 (2022).

24. N. Alessio et al., Low dose radiation induced senescence of human mesenchymal stromal cells and impaired the autophagy process. Oncotarget 6, 8155–8166 (2015).

25. A. Meng, Y. Wang, G. Van Zant, D. Zhou, Ionizing radiation and busulfan induce premature senescence in murine bone marrow hematopoietic cells. Cancer Res 63, 5414–5419 (2003).

26. Y. Zhu et al., Identification of a novel senolytic agent, navitoclax, targeting the Bcl-2 family of anti-apoptotic factors. Aging Cell 15, 428–435 (2016).

27. J. Chang et al., Clearance of senescent cells by ABT263 rejuvenates aged hematopoietic stem cells in mice. Nat Med 22, 78–83 (2016).

28. M. A. Loberg et al., Sequentially inducible mouse models reveal that Npm1 mutation causes malignant transformation of Dnmt3a-mutant clonal hematopoiesis. Leukemia 33, 1635–1649 (2019).

29. J. M. SanMiguel et al., Cell origin-dependent cooperativity of mutant Dnmt3a and Npm1 in clonal hematopoiesis and myeloid malignancy. Blood Adv 6, 3666–3677 (2022).

30. S. Avagyan et al., Resistance to inflammation underlies enhanced fitness in clonal hematopoiesis. Science 374, 768–772 (2021).

31. D. Hormaechea-Agulla et al., Chronic infection drives Dnmt3a-loss-of-function clonal hematopoiesis via IFNγ signaling. Cell Stem Cell 28, 1428–1442.e1426 (2021).

32. S. O. Abegunde, R. Buckstein, R. A. Wells, M. J. Rauh, An inflammatory environment containing TNFalpha favors Tet2-mutant clonal hematopoiesis. Exp Hematol 59, 60–65 (2018).

33. A. G. Bick et al., Inherited causes of clonal haematopoiesis in 97,691 whole genomes. Nature 586, 763–768 (2020).

34. N. Zioni et al., Inflammatory signals from fatty bone marrow support DNMT3A driven clonal hematopoiesis. Nat Commun 14, 2070 (2023).

35. Y. Mei et al., Bone marrow-confined IL-6 signaling mediates the progression of myelodysplastic syndromes to acute myeloid leukemia. J Clin Invest 132, (2022).

36. A. G. Bick et al., Genetic Interleukin 6 Signaling Deficiency Attenuates Cardiovascular Risk in Clonal Hematopoiesis. Circulation 141, 124–131 (2020).

37. W. Liu et al., Blockade of IL-6 signaling alleviates atherosclerosis in Tet2-deficient clonal hematopoiesis. Nat Cardiovasc Res 2, 572–586 (2023).

38. H. Medyouf et al., Myelodysplastic cells in patients reprogram mesenchymal stromal cells to establish a transplantable stem cell niche disease unit. Cell Stem Cell 14, 824–837 (2014).

39. M. H. Raaijmakers et al., Bone progenitor dysfunction induces myelodysplasia and secondary leukaemia. Nature 464, 852–857 (2010).

40. A. Kode et al., Leukaemogenesis induced by an activating beta-catenin mutation in osteoblasts. Nature 506, 240–244 (2014).

41. C. R. Walkley et al., A microenvironment-induced myeloproliferative syndrome caused by retinoic acid receptor gamma deficiency. Cell 129, 1097–1110 (2007).

42. C. R. Walkley, J. M. Shea, N. A. Sims, L. E. Purton, S. H. Orkin, Rb regulates interactions between hematopoietic stem cells and their bone marrow microenvironment. Cell 129, 1081–1095 (2007).

43. I. Aldoss, J. Y. Song, P. T. Curtin, S. J. Forman, Multiple donor-derived leukemias in a recipient of allogeneic hematopoietic cell transplantation for myeloid malignancy. Blood Adv 4, 4798–4801 (2020).

44. S. Chaib, T. Tchkonia, J. L. Kirkland, Cellular senescence and senolytics: the path to the clinic. Nat Med 28, 1556–1568 (2022).

45. H. Fuhrmann-Stroissnigg et al., Identification of HSP90 inhibitors as a novel class of senolytics. Nat Commun 8, 422 (2017).

46. J. N. Farr et al., Targeting cellular senescence prevents age-related bone loss in mice. Nat Med 23, 1072–1079 (2017).

47. A. Kaefer et al., Mechanism-based pharmacokinetic/pharmacodynamic meta-analysis of navitoclax (ABT-263) induced thrombocytopenia. Cancer Chemother Pharmacol 74, 593–602 (2014).

48. P. F. Ruiz-Aparicio, J. P. Vernot, Bone Marrow Aging and the Leukaemia-Induced Senescence of Mesenchymal Stem/Stromal Cells: Exploring Similarities. J Pers Med 12, (2022).

49. E. J. Evans, Jr., J. DeGregori, Cells with Cancer-associated Mutations Overtake Our Tissues as We Age. Aging Cancer 2, 82–97 (2021).

50. S. Srivastava, S. Ghosh, J. Kagan, R. Mazurchuk, The PreCancer Atlas (PCA). Trends Cancer 4, 513–514 (2018).

51. S. Srivastava, S. Ghosh, J. Kagan, R. Mazurchuk, H. I. National Cancer Institute’s, The Making of a PreCancer Atlas: Promises, Challenges, and Opportunities. Trends Cancer 4, 523–536 (2018).

52. A. Ribas, J. D. Wolchok, Cancer immunotherapy using checkpoint blockade. Science 359, 1350–1355 (2018).

53. R. Kuhn, F. Schwenk, M. Aguet, K. Rajewsky, Inducible gene targeting in mice. Science 269, 1427–1429 (1995).

54. R. Gazit et al., Fgd5 identifies hematopoietic stem cells in the murine bone marrow. J Exp Med 211, 1315–1331 (2014).

55. S. Huang et al., An improved protocol for isolation and culture of mesenchymal stem cells from mouse bone marrow. J Orthop Translat 3, 26–33 (2015).

