## Supplementary figures and images for "Mesenchymal Stromal Cell Senescence Induced by *Dnmt3a*-Mutant Hematopoietic Cells is a Targetable Mechanism Driving Clonal Hematopoiesis and Initiation of Hematologic Malignancy"

### Supplemental Figure 1

Supplementary Figure 1.

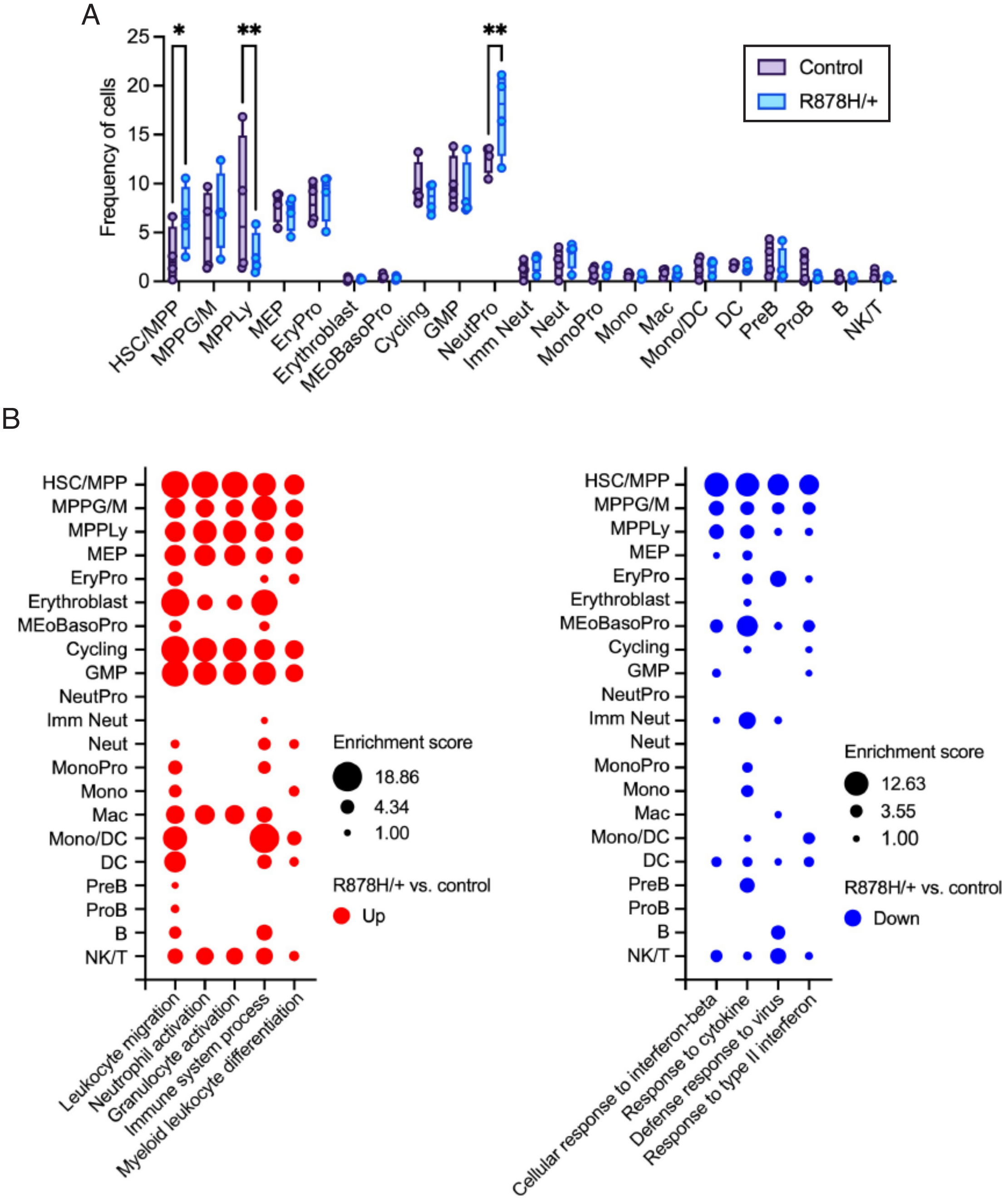

### Supplemental Figure 2

Supplementary Figure 2

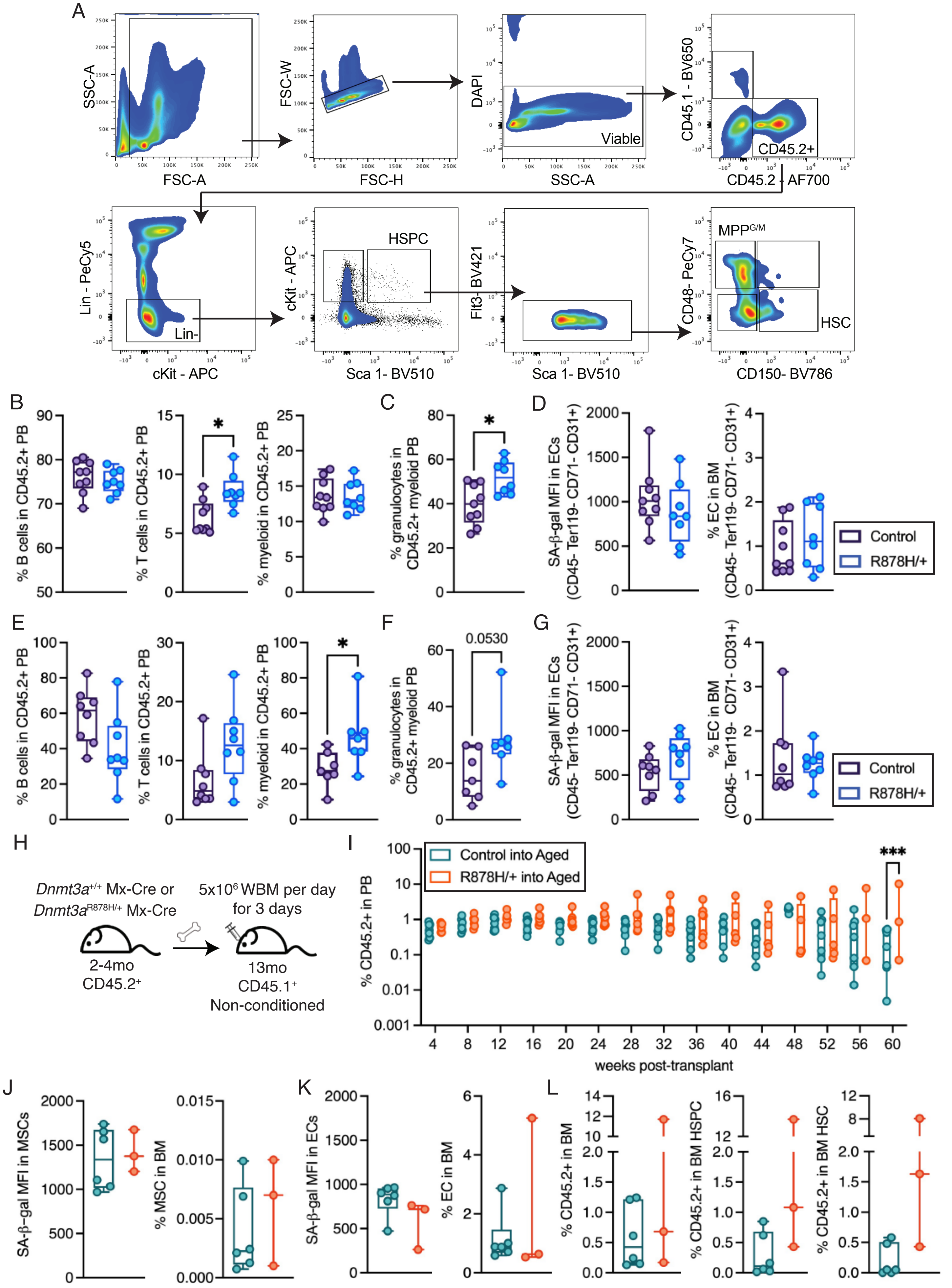

### Supplemental Figure 3

Supplementary Figure 3

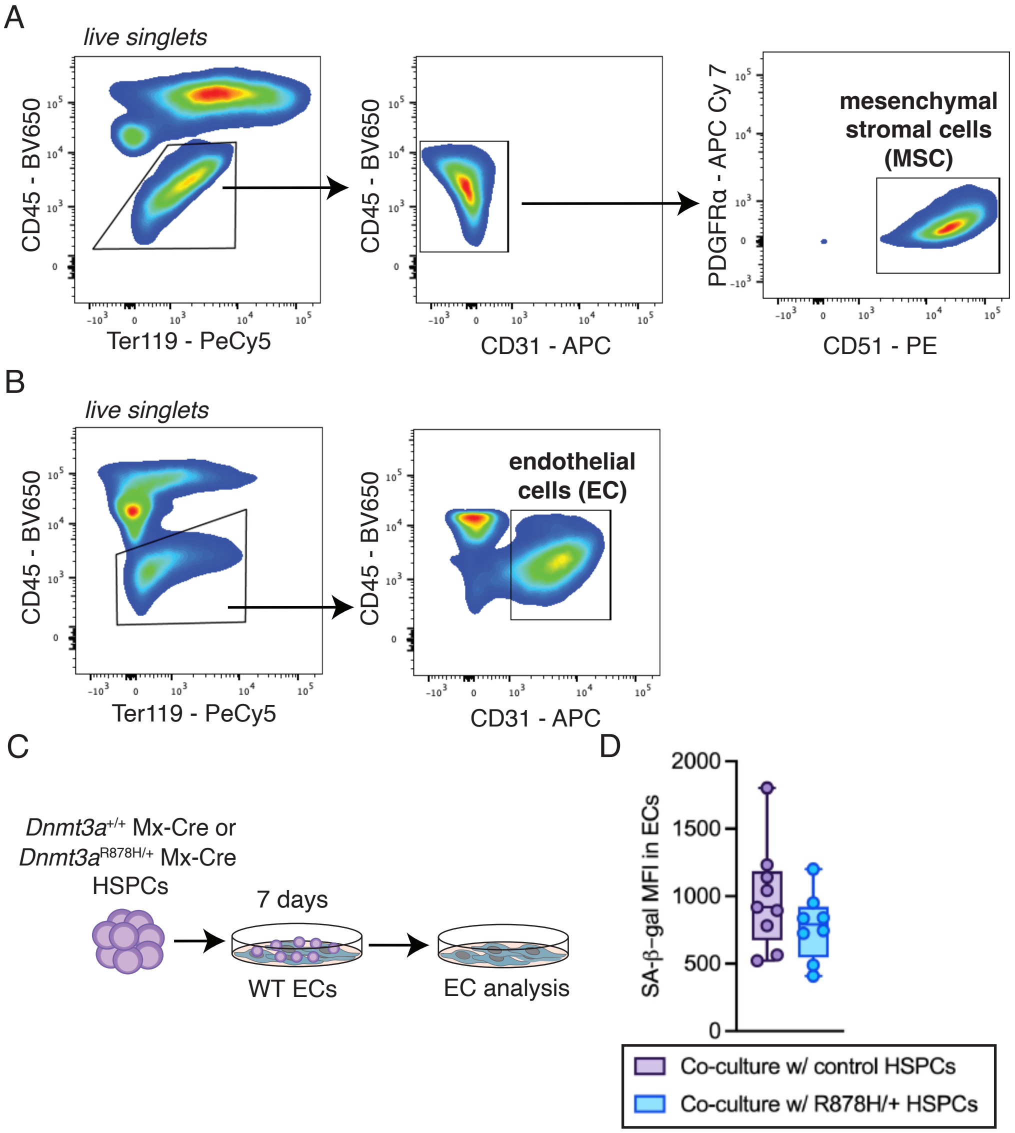

### Supplemental Figure 4

Supplementary Figure 4

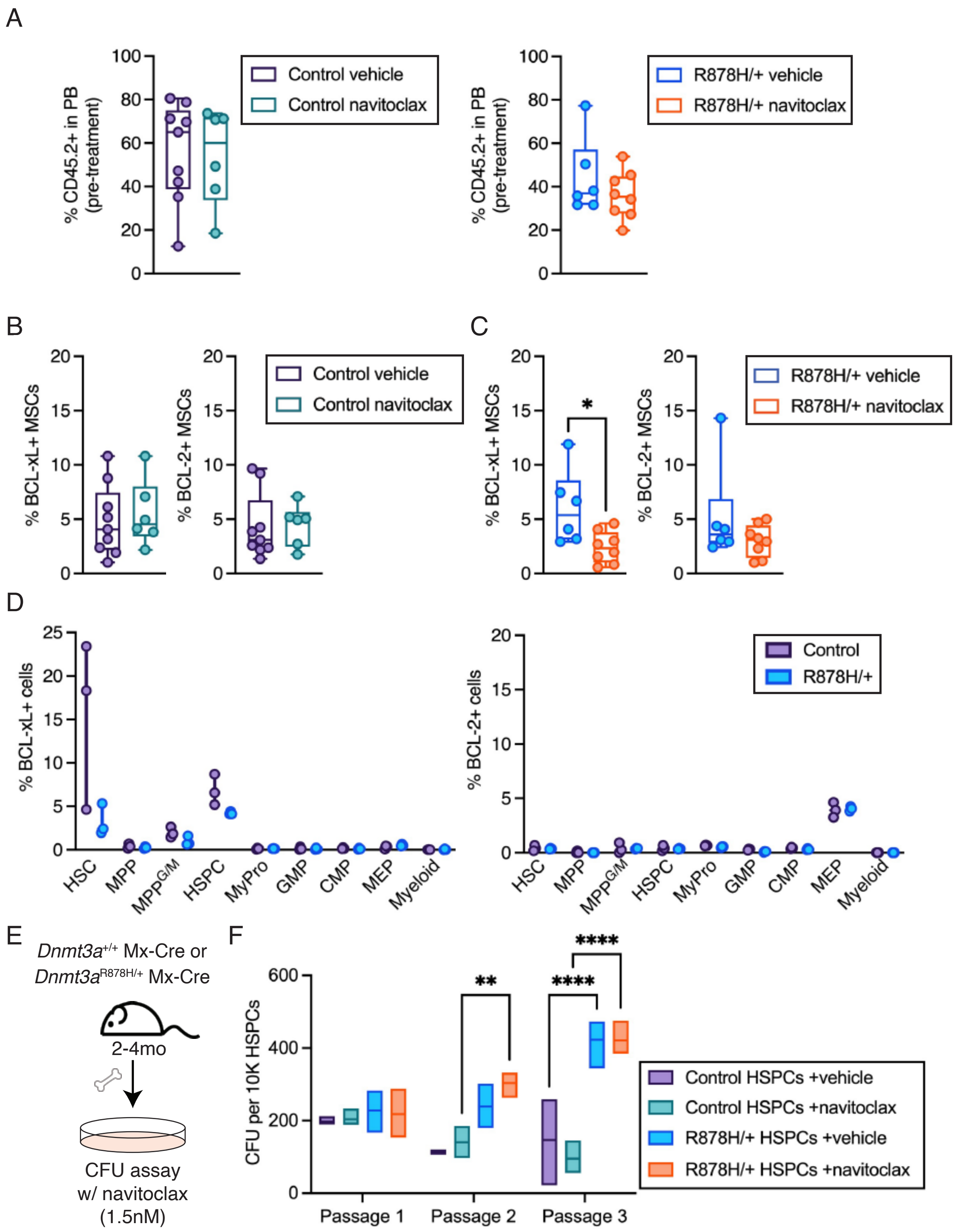
